# Programmable enveloped delivery vehicles for human genome engineering *in vivo*

**DOI:** 10.1101/2022.08.24.505004

**Authors:** Jennifer R. Hamilton, Evelyn Chen, Barbara S. Perez, Cindy R. Sandoval Espinoza, Min Hyung Kang, Marena Trinidad, Jennifer A. Doudna

## Abstract

Viruses and virally-derived particles have the intrinsic capacity to deliver molecules to cells, but the difficulty of readily altering cell-type selectivity has hindered their use for therapeutic delivery. Here we show that cell surface marker recognition by antibody fragments displayed on membrane-derived particles encapsulating CRISPR-Cas9 protein and guide RNA can target genome editing tools to specific cells. These Cas9-packaging enveloped delivery vehicles (Cas9-EDVs), programmed with different displayed antibody fragments, confer genome editing in target cells over bystander cells in mixed cell populations both *ex vivo* and *in vivo*.

This strategy enabled the generation of genome-edited chimeric antigen receptor (CAR) T cells in humanized mice, establishing a new programmable delivery modality with widespread therapeutic utility.

**One-Sentence Summary:** Cell-specific molecular delivery with enveloped delivery vehicles (EDVs) enables genome editing *ex vivo* and *in vivo*.

## Main Text

Therapeutic interventions involving genome editing require the safe and effective delivery of molecules into target cell nuclei (*1*–*3*). Although such capability would transform both clinical and research applications, current non-viral delivery is limited to cells treated *ex vivo (4–6)*, tissues targeted by local administration (*7, 8*), or the liver due to its natural propensity for molecular uptake (*8, 9*). Expansion of *in vivo* genome editing applications will require approaches for molecular delivery to specific cells or organs inside the body following systemic administration.

Retargeting the tropism of viruses or viral vectors is an established delivery strategy involving the surface display of a cell-selective targeting molecule alongside a viral glycoprotein required for cell entry by fusion at the plasma membrane or in the low-pH environment of the endosome (*10*–*13*). Recent progress leverages a mutant form of the vesicular stomatitis virus glycoprotein (VSVG), VSVGmut, that maintains endosomal fusion activity but lacks native LDL receptor binding affinity (*14*–*16*). Pairing VSVGmut with cell-specific targeting molecules can redirect lentiviral transgene delivery and has enabled high-throughput screening of T and B cell receptor libraries to study receptor-antigen interactions (*17, 18*).

Here we show that human cell-specific genome editing can be achieved both *ex vivo* and *in vivo* by pairing the display of VSVGmut with single-chain antibody fragments (scFvs) to create enveloped delivery vehicles that package Cas9 ribonucleoprotein (RNP) complexes (Cas9-EDVs). EDVs use the mechanism of retroviral virus-like particle (VLP) assembly, developed for the transient delivery of Cas9 RNPs (*8, 19*–*25*) or mRNA-encoded genome editors and guide RNAs (*26*–*29*), while avoiding nonspecific editing of bystander cells. We find that Cas9-EDVs achieve targeted genome editing within *in vivo*-generated CAR T cells in mice with a humanized immune system, with no off-target delivery to liver hepatocytes. These data show that EDVs are a programmable platform for delivering molecular cargo to specific cell types for complex genome engineering *in vivo*.

## Results

### Receptor-mediated delivery and genome editing with Cas9-EDVs

A major challenge for *in vivo* delivery of non-viral cargo is the lack of vehicles capable of targeting specific cell types. VLPs can package Cas9 RNP complexes produced by over-expressing Cas9 fused to the C-terminal end of the viral Gag polyprotein during VLP production, but cell-selective VLP targeting has relied on cell infection strategies evolved by enveloped viruses (*20*). To test whether VLPs could be reformulated as programmable EDVs, we first cloned a CD19 targeting antibody as an scFv fused to the stalk and transmembrane domain of CD8α, a strategy commonly used in CAR architecture (*30*) (**Fig. 1A; Fig. S1A, B**). Since Cas9-VLPs bud from the plasma membrane of transfected producer cells, we hypothesized that co-expression of the scFv fusion and VSVGmut together with lentiviral components necessary for Cas9 RNP encapsulation would generate Cas9-EDVs possessing both receptor specificity and endosomal escape capability, respectively.

**Fig. 1.**
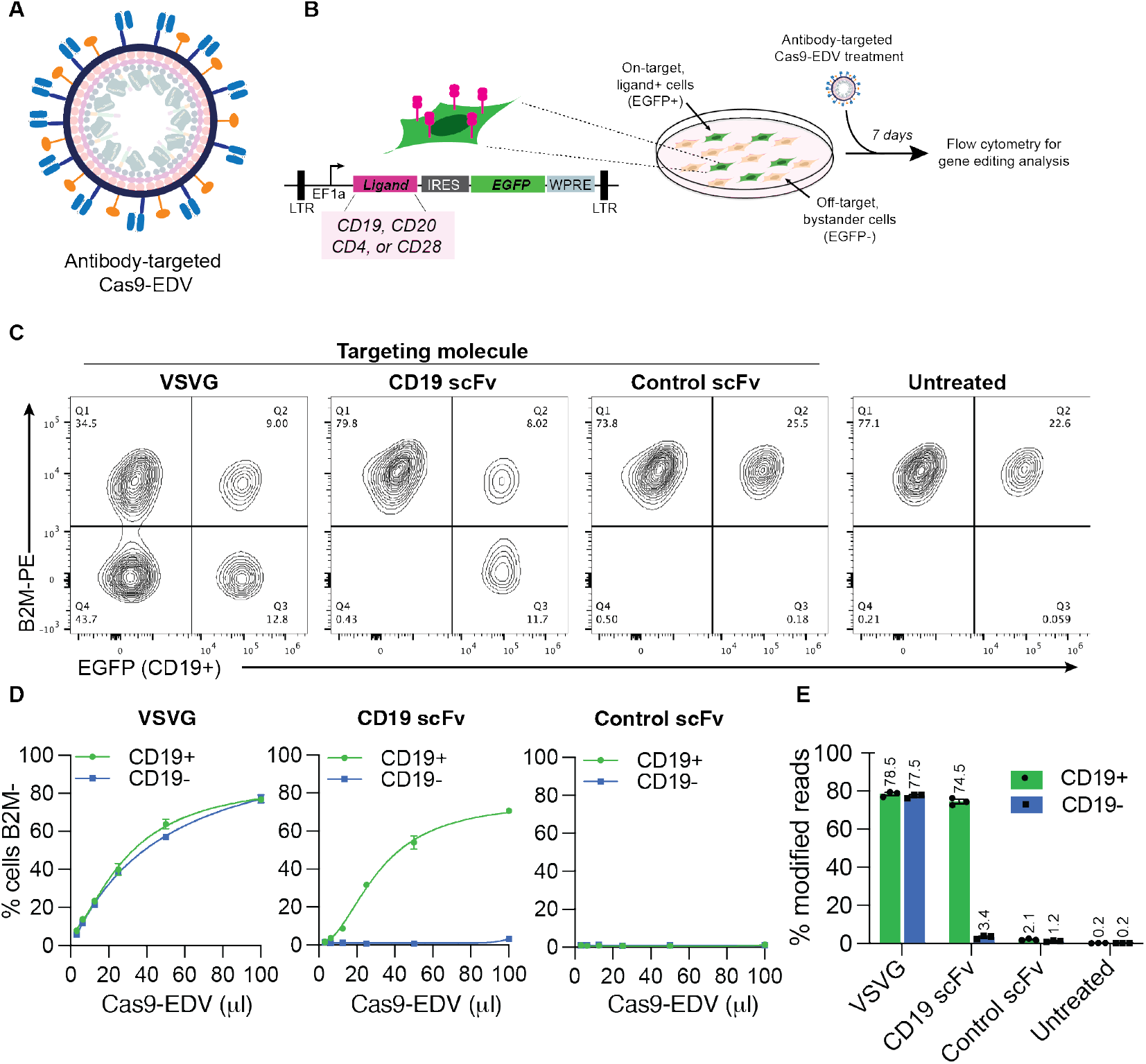
Cell-specific genome editing with antibody-targeted Cas9-EDVs. (**A**) Schematic scFv targeting molecules (blue) and VSVGmut (orange) on the exterior surface of a Cas9-EDV. Cas9-EDVs package pre-formed Cas9-single guide RNA complexes to avoid genetically encoding genome editors within a viral genome. (**B**) Experimental outline and schematic of the lentiviral vector used for engineering HEK293T EGFP cells that express heterologous ligands on the plasma membrane (e.g. CD19). To promote cellular engineering via single lentiviral integration events, engineered cell mixtures were generated via low multiplicity of infection to achieve <25% EGFP+ cells. Engineered cell mixtures were challenged with *B2M*-targeting Cas9-EDVs to test targeting molecule activity. (**C-E**) Assessment of antibody-targeted Cas9-EDV activity. HEK293T and CD19 EGFP HEK293T cells were mixed at an approximate ratio of 3:1 and treated with *B2M*-targeting Cas9-EDVs displaying various targeting molecule pseudotypes. Cas9-EDVs were concentrated 10x, and cells were treated with 50 μl Cas9-EDVs (**C, E**) or in a dilution curve (**D**). Analysis was performed at 7 days post-treatment to assess *B2M* knockout in EGFP+ (on-target) and EGFP- (bystander) cells by flow cytometry (**C, D)** and amplicon sequencing (**E**). N=3 technical replicates for all panels except for the 100 μl dose of CD19-scFv in **(D)** (N = 2). Error bars represent the standard error of the mean.

To analyze the receptor-mediated function of Cas9-EDVs, we generated a HEK293T cell line that co-expresses both the B-cell ligand CD19 and EGFP, enabling assessment of genome editing in on-target, EGFP+ cells, and off-target, EGFP-bystander cells (**Fig. 1B**). We produced Cas9-EDVs containing single guide RNA (sgRNA) targeting the *B2M* gene and outwardly displaying either VSVG, CD19 scFv+VSVGmut, or a control scFv+VSVGmut that should not recognize the target cells in this experiment. In a ∼3:1 mixture of HEK293T and CD19 EGFP 293T cells, the VSVG Cas9-EDVs mediated genome editing in both CD19+ and CD19-populations, whereas the CD19-scFv Cas9-EDVs induced the knockout of *B2M* only in CD19+ cells (**Fig. 1C**). No *B2M* knockout was observed in either the CD19+ or CD19-cells using the control scFv+VSVGmut Cas9-EDVs. Antibody-targeted Cas9-EDV activity was titratable, with up to 74% of target cells exhibiting *B2M* knockout and little to no editing detected in bystander, non-target cells (**Fig. 1D, E**). Antibody-targeted Cas9-EDVs produced genome edits in target cells present at 2-92% of a cell mixture, whereas bystander cell editing was unchanged (**Fig. S1C**). Together these results demonstrate the ability of EDVs to deliver functional molecular cargo in a receptor-mediated fashion.

### Programmable cell-specific genome editing with Cas9-EDVs

Receptor-mediated delivery of genome editing molecules could enable targeted engineering of any cell type as a function of its surface antigens. To test this possibility, we investigated the modularity and programmability of Cas9-EDVs to direct genome editing in HEK293T cells displaying various plasma membrane proteins normally expressed by human immune cells, including CD20, CD4 and CD28 (**Fig. 1B**; **Fig. S2A**). Cas9-EDVs displaying VSVGmut and scFv-based CD20, CD4 and CD28 targeting molecules, generated in both VH-linker-VL and VL-linker-VH orientations (**Table S1**) (*31*), induced up to 80% genome editing that was titratable and selective for ligand(+) over ligand(-) cells (**Fig. S2B**). Not all scFvs produced the same level of on-target cell editing, but in no case was an scFv-displaying EDV able to induce more than minimal editing in off-target cells lacking the cognate surface receptor protein.

Furthermore, in all engineered cell mixtures, ligand(+) and ligand(-) cells were similarly susceptible to genome editing when treated with control Cas9-EDVs that express the VSVG fusogen (**Fig. S2B**). A panel of CD19, CD20 and CD4 antibody-targeted EDVs only mediated genome editing in cells expressing their matched ligand and not in mismatched ligand-expressing cells, demonstrating that delivery requires antibody-antigen interactions (**Fig. S2C**).

### Cas9-EDV optimization enhances genome editing and reveals nonessential components

To enhance Cas9-EDV yield and per-particle editing efficiency ahead of *in vivo* administration, we added p53 nuclear localization signals (NLSs) and nuclear export sequences (NESs) on the C-terminal end of Gag (*8*), improving the editing efficiency of antibody-targeted Cas9-EDVs (**Fig. S3A-D**). Editing efficiency was further increased by expressing sgRNA from both the Gag-NES-NLS-Cas9 and Gag-pol plasmid backbones, as opposed to expression from a separate plasmid (**Fig. 2A, B**). Optimized Cas9-EDVs maintained receptor-mediated delivery specificity except at the highest doses tested (**Fig. S3E**), and Cas9-EDV titration produced a 36-fold enrichment for genome editing on-target cells (79.7%) versus bystander cells (2.2%) (**Fig. S3F**). We also observed improved editing efficiency for optimized Cas9-EDVs versus broadly transducing VSVG Cas9-EDVs when tested on cytokine-stimulated primary human CD34+ cells and cytokine-stimulated and activated primary human T cells *ex vivo* (**Fig. 2C, D**). Surprisingly, optimized Cas9-EDVs mediated genome editing in resting primary human T cells (**Fig. 2E**), which are difficult to edit using standard electroporation approaches.

**Fig. 2.**
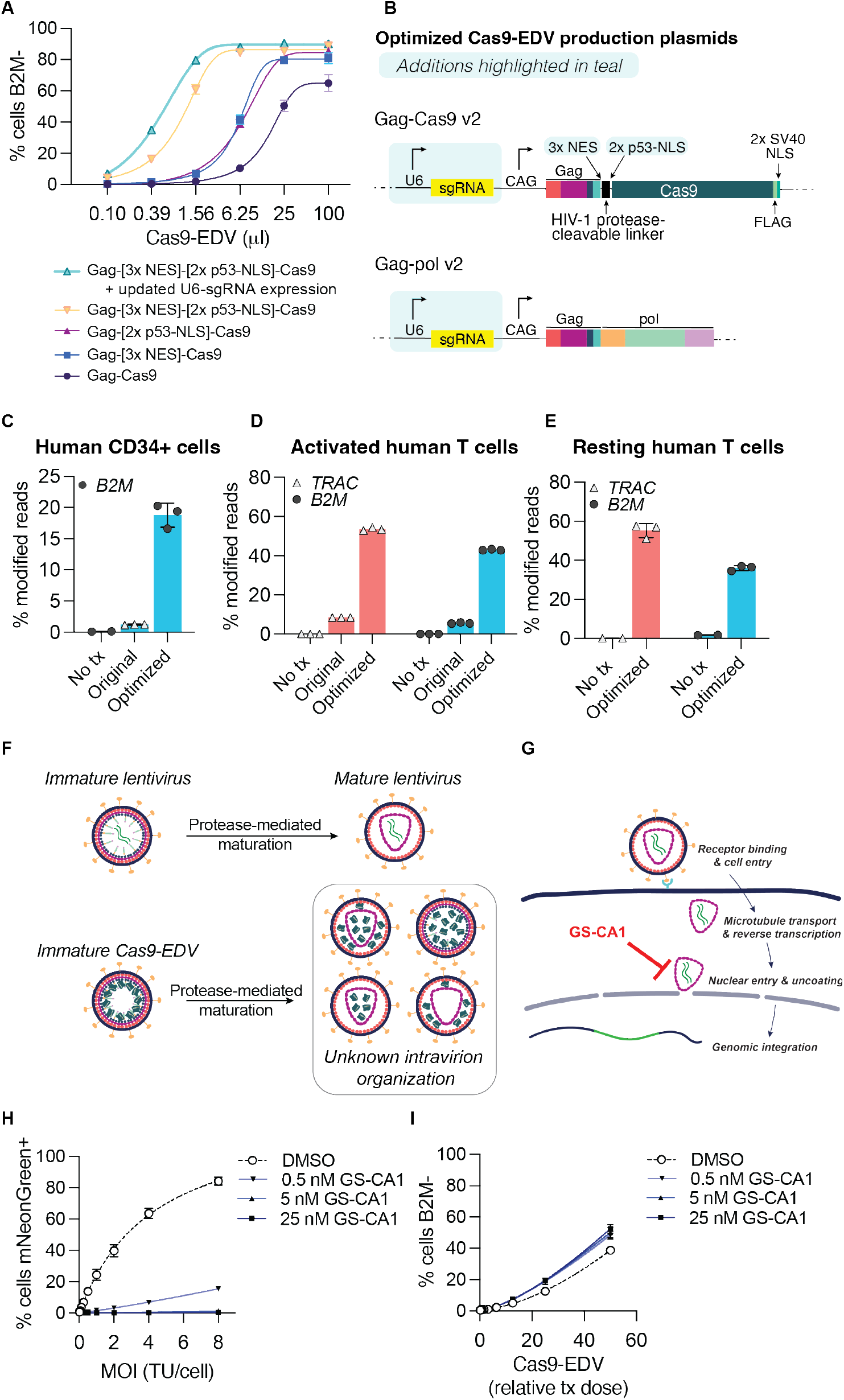
Optimization of Cas9-EDVs for enhanced genome editing activity in primary human cells. (**A**) Genome editing activity comparison of CD19 antibody-targeted Cas9-EDV variants packaging *B2M-*targeted Cas9 ribonucleoproteins (RNPs). Expression of B2M protein was assessed by flow cytometry 7 days post-treatment in CD19-expressing target cells. (**B**) Diagram of the optimized Gag-Cas9 and Gag-pol Cas9-EDV production plasmids; features updated from Hamilton & Tsuchida et al., 2021 are highlighted in teal. (**C-E**) Genome editing activity of optimized VSVG-pseudotyped Cas9-EDVs in primary human CD34+ cells (**C**) and activated (**D**) and resting primary human T cells (**E**). *B2M* or *TRAC g*enome editing was assessed by amplicon sequencing 7 days post-treatment. (**F**) Schematic of potential intra-particle Cas9-EDV configurations for packaged Cas9 RNPs following proteolytic maturation. (**G**) Schematic of the compound GS-CA1 inhibiting either the nuclear import and/or uncoating of an HIV-1 capsid. (**H**) An mNeonGreen lentiviral vector was used to transduce HEK293T cells at the indicated MOI in the presence of GS-CA1 or DMSO. The percent of mNeonGreen-positive cells was assessed by flow cytometry 3 days post-treatment. TU = transducing units. (**I**) *B2M*-targeting Cas9-EDVs, pre-titered such that the highest treatment dose would result in approximately 50% cells B2M-, were used to transduce HEK293T cells in the presence of GS-CA1 or DMSO. B2M expression was assessed by flow cytometry 3 days post-treatment. Error bars represent the standard deviation. N=3 technical replicates were used in all experiments.

We next investigated whether the internal composition of Cas9-EDVs affects editing efficiency, focusing on the lentiviral capsid that forms during proteolytic virion maturation (*32*) (**Fig. 2F**). To probe the role of the capsid in Cas9-EDV delivery, we employed GS-CA1, a small-molecule inhibitor of nuclear import and/or subsequent uncoating of HIV-1 capsid cores (*33, 34*) (**Fig. 2G**). Treatment of target cells with increasing concentrations of GS-CA1 blocked the integration of a lentiviral transgene, which relies on nuclear import from the capsid (**Fig. 2H**), but did not negatively impact the genome editing efficiency of Cas9-EDVs (**Fig. 2I**) or electroporated Cas9 RNPs (**Fig. S3G**). In a separate experiment, we generated a Cas9-EDV variant that relies on the TEV protease to release Cas9 from Gag (“TEVp-Cas9-EDVs,” **Fig. S3H**), preventing the HIV-1 protease-dependent virion maturation required for capsid assembly. Despite the demonstrated loss of Gag proteolytic processing (**Fig. S3I**), TEVp-Cas9-EDVs maintained genome editing activity in treated cells proportional to the amount of Cas9 generated (**Fig. S3J**). These results suggest that the capsid is not required for packaging and delivering Cas9 RNP complexes into target cell nuclei.

### A multiplexed targeting approach for human T cell engineering

Human T cells are important targets for *in vivo* genome engineering applications due to their use in treating cancer and other diseases. Using CD25 expression as a marker, we found that the co-display of CD3 and CD28 targeting molecules on Cas9-EDVs triggered T cell activation and cellular expansion similar to T cells pretreated with commercially-available CD3/CD28 coated magnetic beads (*35*) or engineered lentiviruses (*17*) (**Fig. 3A, B**). CD3+CD28 scFv Cas9-EDV treatment also led to robust levels of genome editing (**Fig. 3C**).

**Fig. 3.**
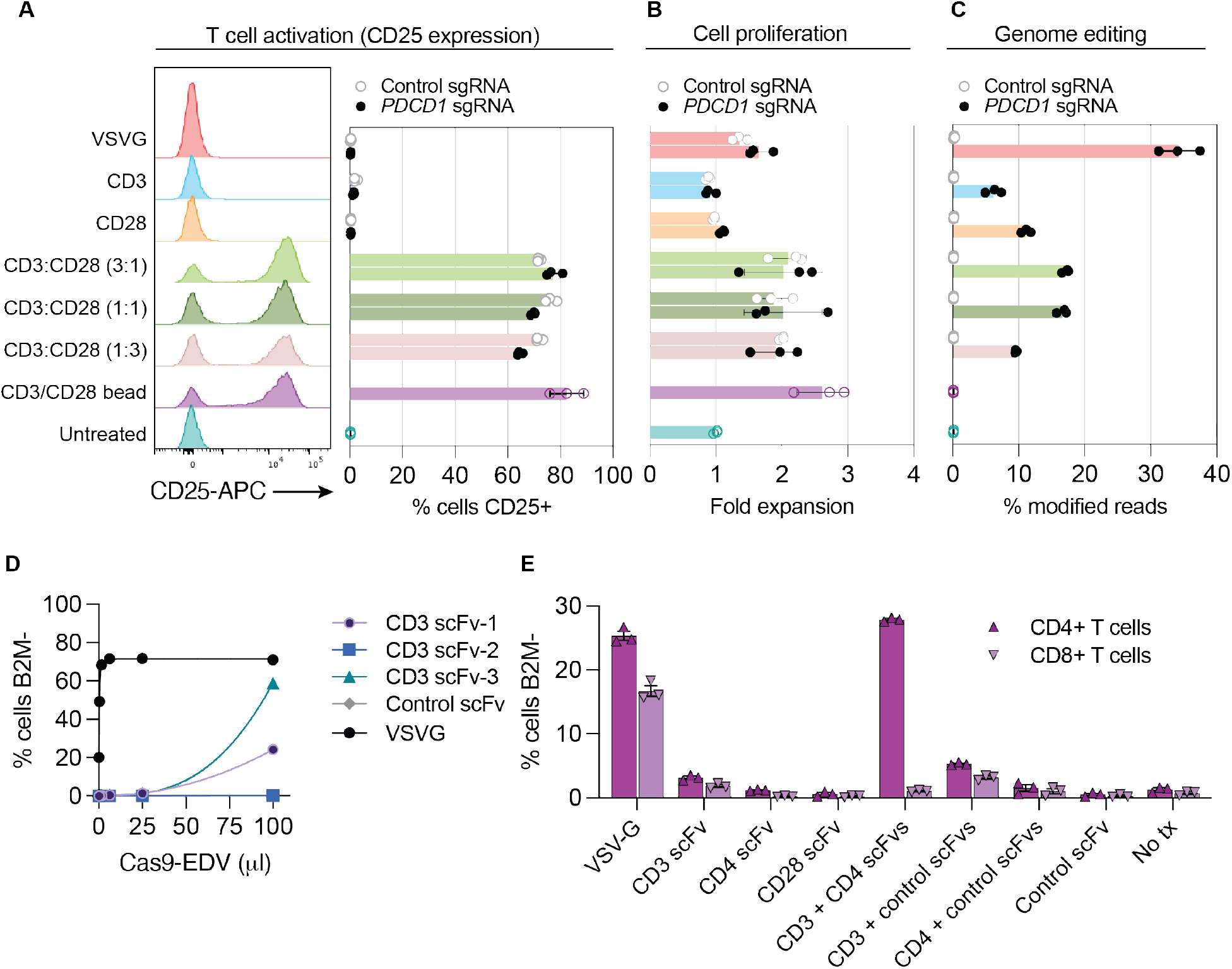
Multiplexed antibody targeting and editing of primary human T cells. (**A-B**) Treating resting human T cells with Cas9-EDVs co-displaying CD3 and CD28 scFvs results in cellular activation (**A**) and proliferation (**B**) as measured by flow cytometry detection of CD25 3 days post-treatment and fold expansion relative to the untreated T cell count, respectively. CD25 expression and cellular proliferation was observed for CD3/CD28 scFv Cas9-EDVs, regardless of whether they packaged Cas9 RNPs targeting *PDCD1* or a non-targeting control. (**C**) Genome editing 3 days post-treatment, as detected by amplicon next-generation sequencing. For A-C, Cas9-EDVs were concentrated 62x and 50 μl was used to treat 30k resting T cells. CD3 scFv-1 and CD28 scFv-2 were tested. (**D**) Screening the mono-display of additional CD3 scFv targeting molecules for *B2M-*targeted Cas9-EDVs on the Jurkat T cell line. B2M expression was assessed by flow cytometry 3 days post-treatment. Cas9-EDVs were concentrated 15x and 50 μl was used to treat 30k Jurkat cells. (**E**) Testing a panel of T cell-targeted, *B2M-*targeting Cas9-EDVs, displaying single or multiplexed scFv targeting molecules. Activated primary human T cells were treated with 1.38×10^8^ Cas9-EDVs displaying one or a combination of CD3 scFv-3, CD4 scFv-2, CD28 scFv-2, and a control scFv, and were assessed for B2M expression in CD4+ and CD8+ T cells by flow cytometry 6 days post-treatment. Error bars represent the standard error of the mean. N=3 technical replicates were for all panels.

Further screening of CD3 and CD45 scFvs revealed additional Cas9-EDV targeting molecules that enabled genome editing of the human Jurkat T cell line (**Fig. 3D; Fig. S4A**). The minimal human T cell editing observed with CD45-targeted Cas9-EDVs may result from a lack of CD45 internalization from the plasma membrane following monoclonal antibody engagement (*36*) (**Fig. S4B, C**). Primary human T cells were susceptible to genome editing using CD3-targeted Cas9-EDVs and, to a lesser extent, CD4-targeted Cas9-EDVs, but not Cas9-EDVs pseudotyped with off-target control scFv targeting molecules (**Fig. 3E; Fig. S4D**). Immunophenotyping of T cells post-Cas9-EDV treatment showed that CD3-targeted Cas9-EDVs direct genome editing in CD4+ and CD8+ subsets of T cells as expected, whereas CD4 scFv-targeted Cas9-EDVs specifically mediated genome editing in the CD4+ subset population (**Fig. 3E; Fig. S4D**). Interestingly, multiplexing CD3 and CD4 scFv targeting molecules on the same Cas9-EDVs led to higher levels of editing than Cas9-EDVs displaying either CD3 or CD4 scFv targeting molecules alone (**Fig. 3E; Fig. S4D**). This observation was antigen-specific, as multiplexing CD3 targeting molecules with off-target control targeting molecules did not enhance genome editing. Receptor cross-linking or aggregation can lead to endocytosis and subsequent lysosomal degradation (*37, 38*), possibly explaining the synergistic increase in genome editing by engaging both CD3 and CD4 receptors.

### T-cell targeted Cas9-EDVs enable complex genome engineering in humanized mice

We investigated the ability of Cas9-EDVs to generate human CAR T cells *in vivo* by delivering both genome editors and transgenes, an advance that could negate the delays and costs associated with current *ex vivo* approaches (*39, 40*). Using immunodeficient mice engrafted with human peripheral blood mononuclear cells (PBMCs) to mimic a humanized immune system, we tested two T cell-targeted vectors for human CAR T cell engineering *in vivo*. One was a lentivirus encoding an α-CD19-4-1BBz CAR-P2A-mCherry transgene to serve as a positive control, and the other was a Cas9-EDV that co-delivers both the lentiviral-encoded CAR and Cas9 RNP complexes to disrupt the T cell receptor alpha constant (*TRAC)* gene (**Fig. 4A)**. Both vectors rely on semi-random integration of a lentivirus for CAR expression, with the Cas9-EDVs additionally packaging Cas9 RNP complexes. Both vectors co-display CD3, CD4 and CD28 scFvs to trigger enhanced cell entry (CD4+CD3) as well as cell activation and proliferation (CD3+CD28).

**Fig. 4.**
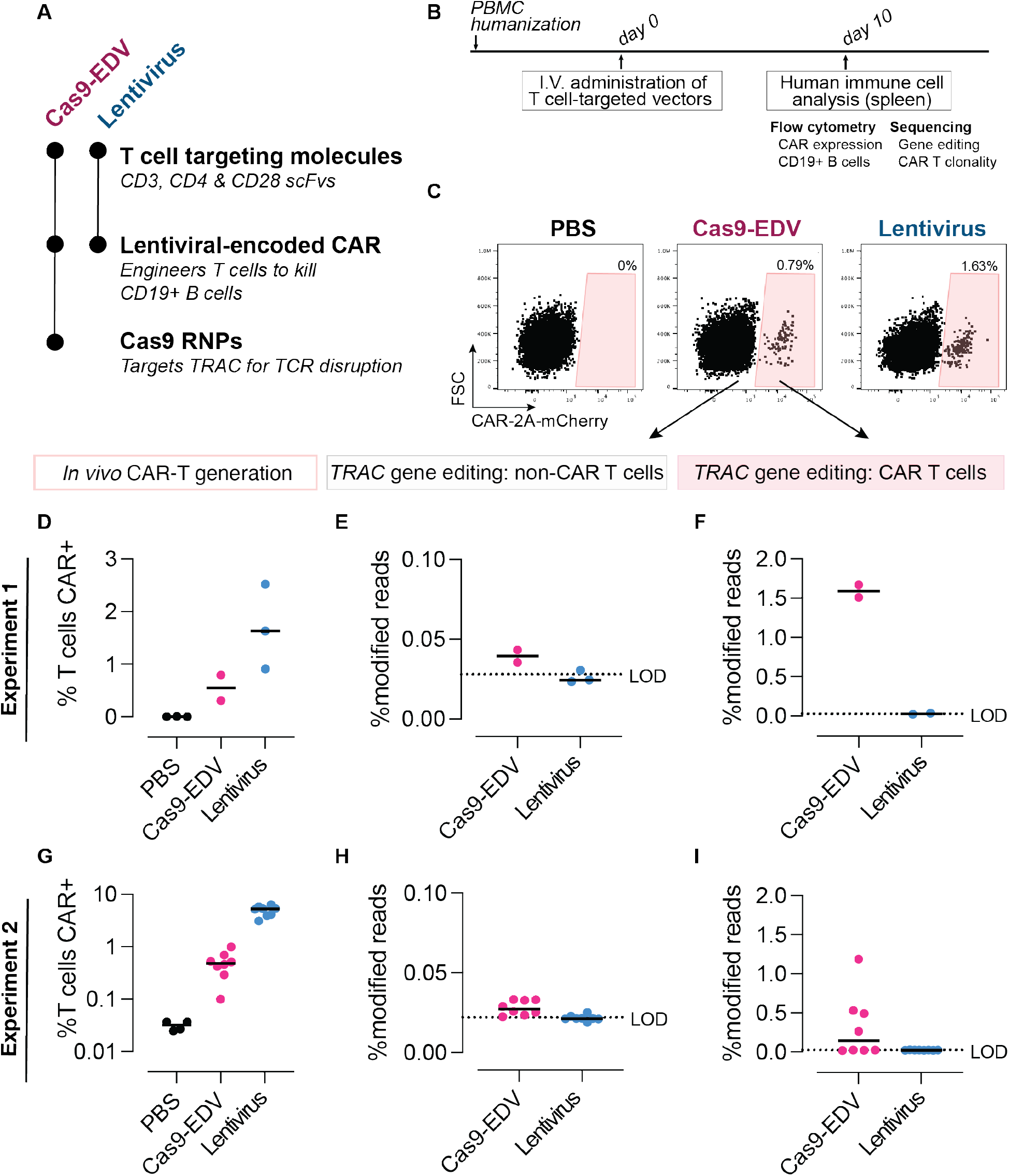
Programmable human cell delivery generates gene-edited CAR T cells *in vivo*. (**A**) Summary of T cell-targeted Cas9-EDVs and lentivirus tested in PBMC-humanized mice. Both particles display multiplexed scFvs (CD3 scFv-3, CD4 scFv-1, and CD28 scFv-2). The Cas9-EDV vector co-packages a lentiviral-encoded CAR-2A-mCherry transgene and Cas9 RNP complexes to disrupt the T cell receptor alpha constant (*TRAC*) gene; the lentivirus encodes the CAR-2A-mCherry transgene. (**B**) Experimental schematic for testing T cell-targeted Cas9-EDVs and lentivirus in PBMC-humanized mice by I.V. (intravenous) retro-orbital injections. (**C**) Representative flow cytometry plots demonstrating that CAR-expressing human T cells are present in the spleens of PBMC-humanized mice 10 days post administration of 1.5×10^9^ Cas9-EDV (N=2) or lentivirus (N=3), quantified in (**D**). PBS = phosphate-buffered saline (**E**) Gene editing is observed in CAR-negative human T cells isolated from mice treated with T cell-targeted Cas9-EDVs, and is enriched in CAR-positive human T cells, **(F)**. One CAR-positive lentivirus sample was excluded in (F) due to failing sequencing. (**G**) CAR-expressing human T cells are detectable in the spleens of PBMC-humanized mice 10 days post administration of 6.2×10^8^ Cas9-EDV (N=8) or lentivirus (N=8) administration. (**H**) Genome editing is observed in CAR-negative human T cells isolated from mice treated with T cell-targeted Cas9-EDVs and is enriched in CAR-positive human T cells, (**I**). For all plots, black lines indicate the median of the data set. LOD = limit of detection, as defined by the average modified reads from lentiviral-treated samples.

T cell-targeting Cas9-EDVs containing the CAR transgene (N=4) or T cell-targeting lentivirus containing the CAR transgene (N=3) were systemically administered and *in vivo* cell engineering was assessed 10 days post-treatment (**Fig. 4B**). CAR-transduced T cells were observed in all mice in which human cells successfully engrafted, as detected by mCherry expression (**Fig. 4C, D; Fig. S5A-C**). In the two Cas9-EDV-treated mice that successfully engrafted with human T cells (N=2 of 4), we observed 1.67% and 1.51% modified alleles in the CAR-transduced T cells, compared to 0.04% and 0.04% in the CAR-negative T cells isolated from the same mice (**Fig. 4E, F**). As expected, no modified alleles were observed in cells isolated from mice treated with the T-cell-targeted lentivirus. Repeating this experiment with mice humanized with PBMCs from a different donor and with more mice per treatment group, we again observed CAR T cells generated *in vivo* in 8/8 mice treated with T cell-targeted Cas9-EDVs and 8/8 mice treated with T cell-targeted lentivirus (**Fig. 4G; Fig. S5D-F**). Again, we observed genome editing only in mice treated with Cas9-EDVs, with higher levels of genome editing in CAR-transduced T cells over CAR-negative T cells (**Fig. 4H, I**). Treatment with the T cell-targeted Cas9-EDV and lentivirus was well tolerated (**Fig. S5G**) and no mCherry+ β-catenin-expressing hepatocytes were observed in the liver (**Fig. S6A, B**). Together, these results indicate that antibody-based targeting of EDVs is a strategy that maintains cell-selective and tissue-specific delivery *in vivo*. Furthermore, these findings suggest that Cas9-EDVs can be used for targeted human cell-type-specific engineering — including both gene insertion and gene disruption — upon single injection *in vivo*.

Because human CD19+ B cells, in addition to T cells, engrafted in the second mouse cohort, we assessed *in vivo* CAR T killing activity. Variable levels of CD19+ B cells were observed in Cas9-EDV-treated mice, and no CD19+ B cells were detected in mice treated with antibody-targeted lentivirus, demonstrating *in vivo* CAR T mediated cytotoxicity (**Fig. 5A; Fig. S7A**). This analysis suggests a model where antibody-derived targeting molecules can direct molecular cargo to specific cells *in vivo* to successfully reprogram cell activity (**Fig. 5B**). Diverse T cell clonotypes were observed for CAR-transduced T cells isolated from mice in both groups (**Fig. 5C**), suggesting that multiple cells were engineered *in vivo*, and did not arise solely through expansion of a single engineered cell. Because clonotype diversity correlated with the number of CAR T cells analyzed (**Fig. S7B**), the clearance of B cells in the lentiviral group was likely attributable to a higher number of CAR T cells generated during the initial *in vivo* transduction. Taken together, these findings offer an approach for generating genome-engineered cells with complex edits that could prove valuable for a wide range of clinical applications in the future.

**Fig. 5.**
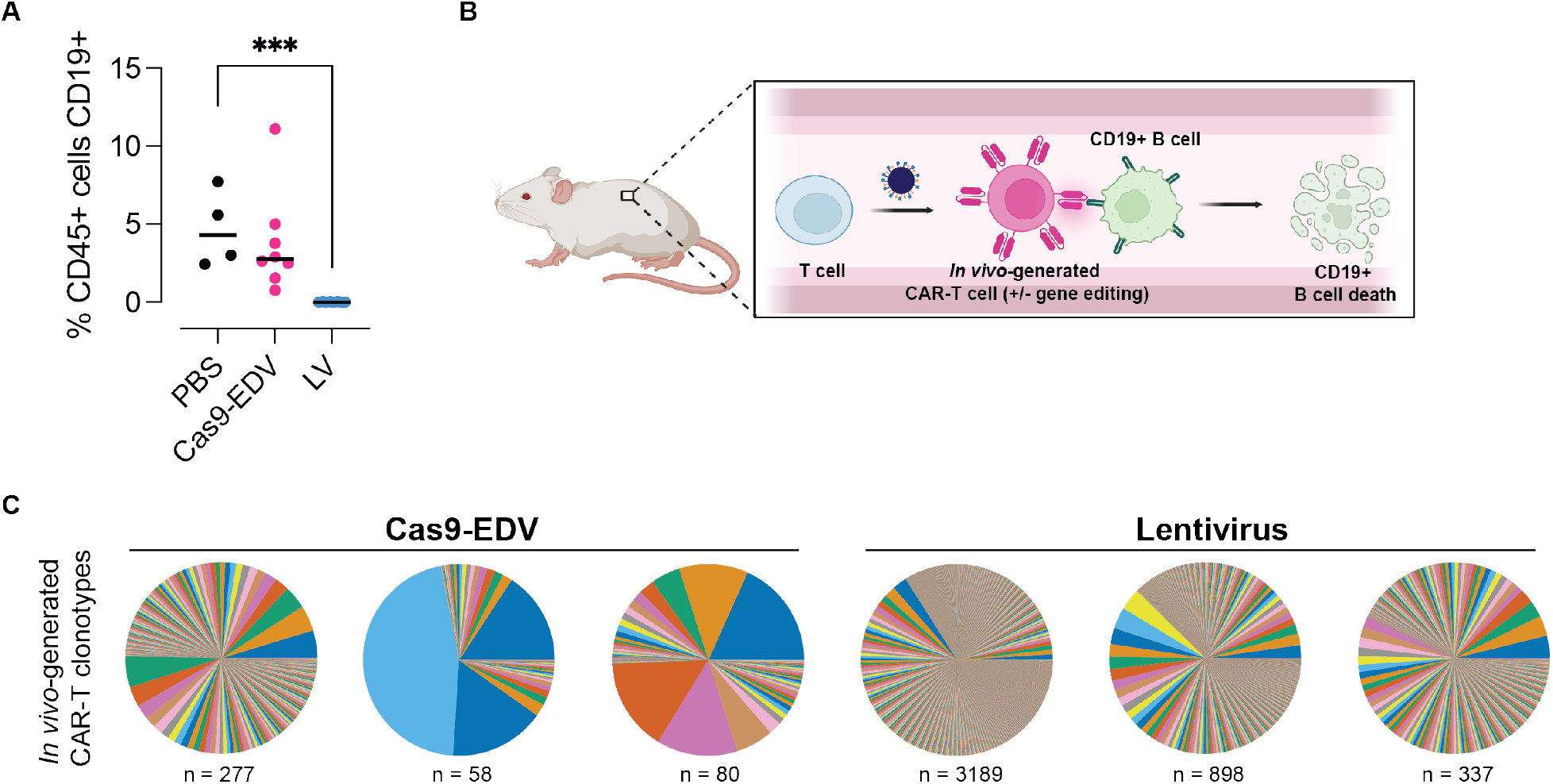
Functional dynamics of cellular engineering *in vivo*. (**A**) Depletion of CD19+ B cells is observed post-administration of T cell-targeted lentivirus (experiment 2). Human CD45+ cells were isolated from PBMC-humanized spleens 10 days post systemic administration, and the percentage of CD19-expressing cells was assessed by flow cytometry. P = *** indicates p<0.001 (unpaired T-test). Black lines indicate the median of the data set. (**B**) Model for the *in vivo* generation of functional CAR-T cells, with or without simultaneous gene editing. Schematic made with Biorender. (**C**) TCR clonotypes of CAR-transduced T cells isolated from humanized mice treated with T cell-targeted Cas9-EDV (mouse numbers 2, 4, 6) or lentivirus (mouse numbers 13, 14, 17) from experiment 2. “n” indicates the unique number of clonotypes detected.

## Discussion

The enveloped delivery vehicles created here combine the molecular packaging and cellular transduction capabilities of a lentivirus with the cell-surface recognition properties of antibodies to deliver Cas9 protein, sgRNAs and transgenes into specific human cell types both *ex vivo* and *in vivo*. Co-display of scFv antibody fragments and the VSVGmut fusogen on the Cas9-EDV envelope provides selectivity of cell transduction. Antibody-directed Cas9-EDVs mediate genome editing in targeted human cells over bystander, non-target cells *in vitro*, and in humanized mice without transducing hepatocytes, thus avoiding a common barrier to selective *in vivo* delivery due to passive liver uptake.

Cas9-EDVs enable complex cell engineering in specific cells, as shown by *in vivo* generation of gene-edited human CAR T cells, with important advantages relative to other *in vivo* delivery methods. First, unlike VLP-mediated delivery (*8*), EDVs can be administered systemically for cell-type specific receptor-mediated delivery of multiple cargo molecules including protein, RNA and DNA. Second, by contrast to viral vector-based methods for delivering DNA-encoded molecules (*41*–*43*), EDVs provide transient delivery of preassembled genome editors whose short lifetime limits off-target editing. In addition, both AAV and lentiviral delivery can involve random transgene integration (*44, 45*) that could be avoided in the future using Cas9 RNP-mediated genome editing for targeted transgene knock-in. Third, distinct from viral vectors (*46*) and lipid nanoparticles (*41*), EDVs do not induce detectable delivery to the liver, which could help avoid toxicity by minimizing the effective concentration necessary for therapeutic benefit. Finally, in contrast to *in vivo* T cell generation using retargeted retroviruses (*47*–*49*), there is no need for T cell activation prior to vector administration because Cas9-EDVs activate T cells during delivery.

One-time, systemic Cas9-EDV administration resulted in ∼0.5% of human T cells that were CAR-positive, in which up to 1.67% modified alleles were observed at the *TRAC* locus in humanized mice. Scaling these efficiencies to the total number of T cells in a typical human (4×10^11^ (*50*)), we would expect to generate 2×10^9^ CAR T cells *in vivo*, of which 3.34×10^7^ CAR T cells would be gene-edited. This compares favorably to the first clinical trial testing CRISPR-Cas9 gene-edited T cells, which infused ∼3.8×10^8^ recombinant TCR T cells, approximately 40% of which were gene-edited at the *TRAC* locus (∼1.5×10^8^) (*4*). Clonotype analysis of *in vivo-* generated CAR T cells indicated that multiple independent transduction events occurred *in vivo*. The incomplete B cell aplasia observed for the T cell-targeted Cas9-EDV dose tested here is likely explained by the number of Cas9-EDV-generated CAR T cells being below a threshold needed for B cell ablation in the first ten days post systemic administration. From the T cell-targeted lentivirus results (5% of T cells *in vivo* became CAR positive), we anticipate that 5% or fewer of all T cells need to undergo *in vivo* engineering to lead to B cell aplasia in this humanized mouse model. Future experiments will assess B cell ablation either at later time points post Cas9-EDV administration to allow more time for cellular expansion, or test higher doses of Cas9-EDVs to generate more CAR T cells initially.

Two aspects of Cas9-EDV composition and cell targeting efficiency were unexpected and warrant further analysis. First, the capsid is not needed, and potentially inhibits Cas9-mediated genome editing, showing that EDV-delivered Cas9 RNP complexes with nuclear localization tags are sufficient to promote nuclear access. Limiting viral structural components to those essential for EDV production may improve the per-particle delivery efficiency of Cas9-EDVs. Second, the finding that not all scFv-based targeting molecules result in equivalent levels of genome editing in target cells suggests that productive EDV delivery requires more than antigen binding. For example, CD45 does not undergo internalization upon antibody binding, which may explain the minimal editing achieved by CD45-scFv Cas9-EDVs (*36, 51*). In addition, differences in scFv-delivery may result from suboptimal targeting molecule display on Cas9-EDVs.

The results reported here merge the single-treatment potential of genome engineering with cell-specific delivery of preassembled genome editors to provide a new approach to selective cell editing *ex vivo* and *in vivo*. These findings also shed light on fundamental aspects of fusion-based cargo delivery that may be further uncovered through the investigation of EDV-based molecular trafficking, offering the potential to use EDVs for fundamental research as well as therapeutic delivery applications.

## Supporting information

Table S1. Information on single-chain variable fragment targeting molecule sequences

## Acknowledgments

We thank all members of the Doudna laboratory for their thoughtful input on this project, particularly Connor Tsuchida and Abby Stahl. We thank Netra Krishnappa and the IGI NGS Core for assistance with next-generation sequencing and Linda Vo for help with human CD34+ cell experiments. Jamie Cate, Matthew Kan, Enrique Lin Shiao, Kevin Wasko, Abdullah Syed, and Ross Wilson provided helpful comments on the manuscript. We also thank Wes Sundquist for helpful discussions and the suggestion to use GS-CA1 for capsid disruption, and Stephen Yant (Gilead Sciences, Inc.) for providing GS-CA1.

## Funding

Centers for Excellence in Genomic Science of the National Institutes of Health, grant RM1HG009490 (JAD)

Somatic Cell Genome Editing Program of the Common Fund of the National Institutes of Health, grant U01AI142817-02) (JAD)

Howard Hughes Medical Institute (JAD)

National Institute of General Medical Sciences of the National Institutes of Health, grant1K99GM143461-01A1 (JRH)

Jane Coffin Childs Memorial Fund for Medical Research. (JRH)

## Author contributions

Conceptualization: JRH, JAD

Methodology: JRH, EC, BSP, CSE, JAD

Software: MT

Investigation: JRH, EC, BSP, CSE, MHK

Visualization: JRH, EC, BSP, MHK, MT

Funding acquisition: JRH, JAD

Project administration: JRH, JAD

Writing – original draft: JRH, EC, BSP, JAD

Writing – review & editing: JRH, EC, BSP, CSE, MHK, MT, JAD

