## Supplementary material for "Programmable enveloped delivery vehicles for human genome engineering *in vivo*": Methods and Supplementary Materials

### Materials and Methods

Plasmid construction

VSVGmut (K47A R354A VSVG) sequence was human codon-optimized and synthesized as a gBlock (Integrated DNA Technologies) and cloned into the pCAGGS expression plasmid. To generate the CD19 scFv-1 expression plasmid, the sequence encoding the CD8α signal peptide, myc epitope tag, scFv, and CD8α stalk and transmembrane domain of α-CD19-4-1BBζ-P2A-mCherry (*20*, *52*, *53*) was subcloned into pCAGGS. This plasmid was subsequently used as an entry plasmid for cloning all other scFv antibody fragments: the CD8α signal peptide, myc tag, and scFv sequences were dropped out by EcoRI/Esp3I restriction digest (New England Biolabs) and new DNA sequences encoding CD8α signal peptide and scFv were inserted. This cloning strategy resulted in removing the n-terminal myc epitope tag and adding a serine amino acid residue between the scFv and CD8α hinge domains. A flexible linker (GGGGSGGGGSGGGGSS) was used to link VH and VL domains of source monoclonal antibody sequences. If the antibody source sequence was already an scFv, the linker from the source sequence was used. Except for CD19 scFv-1, all antibody fragment sequences were human codon-optimized and synthesized as eBlock Gene Fragments (Integrated DNA Technologies). InFusion cloning (Takara Bio) was used to generate all plasmids. Additional information on the scFv targeting molecules and source sequences can be found in **Table S1**.

A second-generation lentiviral transfer plasmid encoding expression of EF1α promoter - CAR-P2A-mCherry (*20*) was digested with XbaI and MluI (New England Biolabs) to drop out the CAR-P2A-mCherry transgene. Human CD19 (Uniprot #Q71UW0) DNA was ordered as a gBlock (Integrated DNA Technologies) and IRES-EGFP (amplified from the Xlone TRE3G MCS-TEV-Halo-3XF IRES EGFP-Nuc-Puro plasmid, a gift from the Darzacq/Tijan Lab) sequences were inserted using InFusion cloning (Takara Bio). This cloning strategy inserted a MluI restriction digest site 3' of the CD19 stop codon and removed the MluI restriction digest site 3' of the EGFP stop codon. Human CD4 (Uniprot #P01730), CD20 (Uniprot #P11836), and CD28 (Uniprot #P10747) amino acid sequences were human codon-optimized for synthesis and ordered as an eBlock (CD28) or gBlocks (CD20, CD4) (Integrated DNA Technologies). Ligand-encoding sequences were cloned by restriction digest removal of CD19-encoding sequence from the EF1α-CD19 IRES-EGFP lentiviral plasmid using XbaI & MluI (New England Biolabs) and inserted with InFusion cloning (Takara Bio). VSVGmut and scFv targeting plasmids were prepared using the HiSpeed Plasmid Maxi or Plasmid Plus Midi kits (QIAGEN). Lentiviral plasmids were prepared with the QIAprep Spin Miniprep Kit (QIAGEN). All plasmids were sequence-confirmed (UC Berkeley DNA Sequencing Facility, Quintara Bio, or Primordium Labs) before use.

The p53-NLS aa 305-322 sequence was obtained by reverse transcription PCR using RNA extracted from Raji cells as a template with the SuperScript™ III One-Step RT-PCR System with Platinum™ Taq DNA Polymerase (Thermo Fisher). 2x-p53 NLS was constructed by linking two p53 NLS sequences by a flexible linker (GGSGG) and the 2x-p53 NLS sequence was inserted into Gag-Cas9 (Addgene plasmid #171060) with InFusion cloning (Takara Bio). Gag-Cas9 and Gag-[2x p53-NLS]-Cas9 were digested with MfeI-HF and AgeI-HF (New England Biolabs). 3x NES sequence was human codon-optimized, synthesized as a gBlock (Integrated DNA Technologies), and inserted with InFusion cloning (Takara Bio) to generate Gag-[3x NES]-[2x p53-NLS]-Cas9 .

The U6-sgRNA expression cassette was cloned into the plasmid backbone of psPax2 as follows: Gag-[3x NES]-[2x p53-NLS]-Cas9 was digested with SalI-HF (New England Biolabs). The Gag-pol expression plasmid psPax2 (Addgene plasmid #12260) was first digested with AflII and SacI (New England Biolabs) to remove a SalI restriction site. The modified psPax2 was then digested with SalI-HF (New England Biolabs). The U6-sgRNA expression cassette was amplified from the spyCas9 sgRNA-BsmBI-Destination plasmid (Addgene plasmid #171625) and inserted into the digested Gag-[3x NES]-[2x p53-NLS]-Cas9 and psPax2 with InFusion cloning (Takara Bio). Oligos encoding guide RNA spacers (*B2M*: 5'- GAGTAGCGCGAGCACAGCTA; *TRAC*: 5'-AGAGTCTCTCAGCTGGTACA; *PDCD-1*: 5'-CGACTGGCCAGGGCGCCTGT; *tdTomato*: 5'-AAGTAAAACCTCTACAAATG; control: 5'-GTATTACTGATATTGGTGGG) were ordered from IDT, phosphorylated, annealed and ligated into BsmBI-digested sgRNA expression vectors.

Tissue culture and cell line generation

Lenti-X and HEK293T cells, obtained and authenticated by the UC Berkeley Cell Culture Facility, were cultured in DMEM (Corning) supplemented with 10% fetal bovine serum (VWR) and 100 U/ml penicillin-streptomycin (Gibco) ("cDMEM"). To generate lentiviruses encoding EF1α-ligand IRES-EGFP, 3.5-4 million Lenti-X cells were plated in a 10 cm tissue culture dish (Corning) and transfected with 1 µg pCMV-VSV-G (Addgene plasmid #8454), 10 µg psPax2 (Addgene plasmid #12260), and 10 µg of EF1α-ligand IRES-EGFP lentiviral transfer plasmid using polyethylenimine (Polysciences Inc.) at a 3:1 PEI:plasmid ratio. Lentiviral-containing supernatants were harvested two days post-transfection and passed through a 0.45 µm PES syringe filter (VWR). Ligand-expressing cells were generated by transducing HEK293T cells (100,000 per well in a 12-well dish) with lentivirus (0.15-1 ml) in a total well volume of 1 ml. Four days post-transduction, flow cytometry was used to identify cell mixtures where <25% of cells were expressing EGFP. Following expansion, CD19 EGFP HEK293T cells were additionally sorted for EGFP expression using an SH800S cell sorter (Sony Biotechnology) to generate a population of cells ~100% CD19+EGFP+ cells.

T cell culture

Cryopreserved human peripheral blood mononuclear cells (PBMCs, AllCells) were thawed in X-VIVO 15 (Lonza) with 50 μM 2-Mercaptoethanol (Gibco), 5% fetal bovine serum (VWR), and 10 mM N-acetyl L-cysteine (Sigma-Aldrich). This media was also used to culture the T cells isolated from PBMCs. To isolate T cells, PBMCs were incubated in 100 μg/mL DNase I solution (Stem Cell Technologies) at room temperature for 15 minutes and resuspended in EasySep™ Buffer (Stem Cell Technologies). Aggregated suspensions were filtered through a 37 μm cell strainer (Stem Cell Technologies) and incubated with EasySep™ Human T Cell Isolation Cocktail (Stem Cell Technologies). Then, EasySep™ Dextran RapidSpheres™ (Stem Cell Technologies) were used to separate T cells via an EasySep™ Magnet (Stem Cell Technologies). Dynabeads™ Human T-Activator CD3/CD28 (Gibco) and recombinant human 500 U/mL IL-2 (Peprotech), 5 ng/mL IL-7 (Peprotech), and 5 ng/mL IL-15 (R&D Systems) were used to stimulate and activate T cells for two days before treatment. Pre-activated T cells were cultured in media containing 500 U/mL IL-2 (Peprotech).

CD34+ cell culture

Cryopreserved G-CSF-mobilized human CD34+ cells were acquired from AllCells. Cells were thawed, resuspended inIMDM (Gibco) spun, and then were cultured in StemSpan™ SFEM II media (Stem Cell Technologies) with 100 U/ml penicillin-streptomycin (Gibco) and the cytokine cocktail StemSpan™ CC110 (Stem Cell Technologies). Cells were treated with 8 μM cyclosporine H (Sigma-Aldrich) for 24 hours before treatment. Additionally, cells were treated with 1 μg/ml poloxamer (BASF) at the time of treatment. Cas9-EDVs were concentrated 50-fold using Lenti-X Concentrator (Takara Bio), resuspended in SFEM II media, and mixed with 40k CD34+ cells in a final well volume of 100 μl. Transduction was performed in a U-bottom 96-well plate for 24 hours on an orbital shaker before cells were spun, expanded 1:1, and cultured stationary until analysis.

Cas9-EDV production

Cas9-EDVs (formerly known as "Cas9-VLPs") were produced as previously described (*20*). Briefly, VSVG-pseudotyped Cas9-EDVs were produced by seeding 3.5-4 million Lenti-X cells (Takara Bio) into 10 cm tissue culture dishes (Corning) and transfecting the next day with 1 μg pCMV-VSV-G (Addgene plasmid #8454), 6.7 μg Gag-Cas9 (Addgene plasmid #171060), 3.3 μg psPax2 (Addgene plasmid #12260), and 10 μg U6-sgRNA (Addgene plasmid #171635 or Addgene plasmid #171634) using polyethylenimine (Polysciences Inc.) at a 3:1 PEI:plasmid ratio. Antibody-targeted Cas9-EDVs were produced in the same way as VSVG Cas9-EDVs, except that the pCMV-VSV-G plasmid was omitted, and 7.5 μg of scFv targeting plasmid and 2.5 μg of VSVGmut were included during transfection. For scFv multiplexing, a total of 7.5 μg scFv plasmids was used, split 1:1 or 1:1:1, unless otherwise described in the figure legend. For Cas9-EDVs used to treat primary human cells (T cells, CD34+ cells) or humanized mice, media was changed 6-18 hours post transfection into Opti-MEM (Gibco). Two days post-transfection (or media change), Cas9-EDV-containing supernatants were harvested, passed through a 0.45 μm PES syringe filter (VWR), and concentrated with Lenti-X Concentrator (Takara Bio) according to the manufacturer's instructions. Concentrated Cas9-EDVs were resuspended in Opti-MEM (Gibco) at a final concentration of 10x unless otherwise noted in the figure legend.

Cas9-EDVs were stored at 4°C or frozen at -80°C within an isopropanol-filled freezing container until use.

To optimize Cas9-EDVs, variants with different Gag-Cas9 polypeptides were produced in the same way as above, except that 6.7 µg of each Gag-Cas9 variant (Gag-[3x NES]-Cas9, Gag-[2x p53-NLS]-Cas9, and Gag-[3x NES]-[2x p53-NLS]-Cas9) was used instead of Gag-Cas9 during transfection. Gag-[3x NES]-[2x p53-NLS]-Cas9 EDVs with U6-sgRNA expression from plasmid backbones were produced similarly, except that the U6-B2M plasmid was omitted, and 6.7 µg of U6-sgRNA Gag-[3x NES]-[2x p53-NLS]-Cas9 and 3.3 µg of U6-sgRNA psPax2 were included instead of the Gag-Cas9 and psPax2 plasmids during transfection. To capture both improvements in Cas9-EDV particle production and the per-particle editing efficiency of Cas9 EDVs, we produced equivalent numbers of transfected 10 cm tissue culture dishes (Corning) of each Cas9-EDV variant tested.

For humanized mouse experiments, Cas9-EDVs were produced by transfecting Lenti-X cells with 2.5 µg of each scFv targeting plasmid (CD3 scFv-3, CD4 scFv-1, and CD28 scFv-2), 2.5 µg of VSVGmut, 3.3 µg of U6-TRAC Gag-[3x NES]-[2x p53-NLS]-Cas9, 6.7 µg of U6-TRAC psPax2, and 2.5 µg of the lentiviral transfer plasmid encoding an α-CD19-4-1BBz CAR-P2A-mCherry transgene, as optimized in (*20*). Lentivirus was produced in the same way, except that U6-TRAC Gag-[3x NES]-[2x p53-NLS]-Cas9 and U6-TRAC psPax2 were omitted, and 10 µg psPax2 was included. Cas9-EDV- and LV-containing supernatants were passed through a 0.45 µm PES filter bottle (Thermo Fisher) and concentrated via ultracentrifugation by floating the supernatant on top of a cushioning buffer of 30% (w/v) sucrose in 100 mM NaCl, 10 mM Tris-HCl pH 7.5, 1 mM EDTA pH 8.0, at 25,000 rpm with a SW28 Ti rotor (Beckman Coulter) for 2 hours at 4°C in polypropylene tubes (Beckman Coulter). After ultracentrifugation, the Cas9-EDV pellet was resuspended in sterile Dulbecco's phosphate-buffered saline (DPBS) (Gibco).

Cas9-EDV titer quantification

The QuickTiter™ Lentivirus Titer Kit (Lentivirus-Associated HIV p24) (Cell Biolabs, INC) was used to quantify Cas9-EDV particle number. Cas9-EDVs were diluted 1:1,00-100,000 and the ELISA was performed according to the manufacturer's directions. 450 nm absorbance was measured plate reader (BioTek). Cas9-EDV p24 content was calculated by comparison to serial dilution of a p24 standard and guidance from the manufacturer was used to convert p24 quantity into particle number (Cell Biolabs, INC).

Western blot analysis

Cas9-EDVs were mixed with Laemmli buffer containing 10% 2-mercaptoethanol and heating at 90C for 5 minutes. Proteins from whole cell lysates were separated by 4%–20% SDS-PAGE gel (Bio-Rad) and transferred to an Immun-Blot® low fluorescence PVDF membrane (Bio-Rad). Membranes were blocked and incubated with primary antibodies at 4 °C overnight followed by secondary antibodies at room temperature for 1 hour. Primary antibodies were anti-FLAG M2 (F1804, RRID:AB_262044, Sigma-Aldrich) and anti-HIV1 p24 (ab9071, RRID:AB_306981, abcam). Secondary antibodies were anti-mouse Alexa Fluor™ 488 (A28175, RRID:AB_2536161, Thermo Fisher) and anti-rabbit Alexa Fluor™ 647 (A-31573, RRID:AB_2536183, Thermo Fisher). Imaging was performed using the Odyssey imaging system (LI-COR).

GS-CA1 experiments

To generate lentiviruses encoding EF1α-mNeonGreen, 3.5-4 million Lenti-X cells were plated in a 10 cm tissue culture dish (Corning) and transfected with 1 µg pCMV-VSV-G (Addgene plasmid #8454), 10 µg psPax2 (Addgene plasmid #12260), and 2.5 µg of EF1α-mNeonGreen lentiviral transfer plasmid using polyethylenimine (Polysciences Inc.) at a 3:1 PEI:plasmid ratio. Lentiviral-containing supernatants were harvested two days post-transfection and passed through a 0.45 µm PES syringe filter (VWR). Lentiviral supernatants were concentrated 20x with Lenti-X Concentrator (Takara Bio) according to the manufacturer's instructions, resuspended in Opti-MEM (Gibco), aliquoted, and frozen at -80°C for future use. *B2M*-targeted VSVG Cas9-EDVs were produced, as described above. The mNeonGreen lentivirus stock was pre-titered on HEK293T cells: Cas9-EDV and mNeonGreen lentivirus samples were diluted in Opti-MEM in a 2-fold dilution series. 50 µl of each dilution series was mixed with 15,000 HEK293T cells in 50 µl cDMEM in triplicate in a 96-well plate. For the lentiviral sample, the percentage of mNeonGreen-positive cells was assessed by flow cytometry three days post-transduction. Wells where the percent of mNeonGreen+ cells was ≤25% were used to calculate the transducing units (TU) per ml. The genome editing activity of the Cas9-EDV stock was pre-titered similarly, except that B2M expression was assessed by flow cytometry at three days post-treatment to calculate Cas9-EDV volume that resulted in approximately 50% cells negative for B2M expression.

The HIV-1 capsid inhibitor GS-CA1 (Gilead Sciences, Inc.) was diluted to a working stock of 100 µM using DMSO. 15,000 HEK293T cells were transduced with mNeonGreen lentivirus or *B2M*-targeting Cas9-EDVs and simultaneously treated with 0, 0.5, 5, or 25 nM GS-CA1 in a total well volume of 100 µl (0.05% DMSO final). mNeonGreen and B2M expression were assessed by flow cytometry three days post-treatment. As a control, *B2M* sgRNA and Cas9 protein (Integrated DNA Technologies) were complexed at a 2:1 ratio for 15 minutes at 37°C, and 50 pmol Cas9 RNPs were nucleofected into 200,000 HEK293T cell using the SF Cell Line 4D-Nucleofector Kit and 4D-Nucleofector instrument (Lonza) using pulse code CM-130. Post nucleofection, cells were brought up to 100 µl with pre-warmed cDMEM and incubated at 37°C for 15 minutes. Nucleofected cells were plated at 15,000 per well of a 96-well plate and treated with 25 nM GS-CA1, 0.05% DMSO, or Opti-MEM. B2M expression was assessed by flow cytometry three days post-nucleofection.

Humanized mouse experiments

All studies and procedures were in accordance with the established NIH guidelines for animal care and use and were approved by the University of California, Berkeley, Animal Care and Use Committee (ACUC). All experimental and control animals were housed under the same conditions as approved by the Berkeley Office of Laboratory Animal Care (OLAC). Human peripheral blood mononuclear cell-engrafted NSG™ mice (745557) were purchased from Jackson Laboratory. "Experiment 1" and "experiment 2" mouse cohors were engrafted with cells from unique human donors. Following anesthesia induction with 2-3% isofluorane, 100 µl of Cas9-EDVs, lentivirus, or phosphate-buffered saline (PBS) was administered by retro-orbital injection. 10 days post-treatment, mice were euthanized with CO2 to harvest spleen and liver. See "Immunofluorescent staining and imaging" for downstream liver sample processing. Single-cell suspensions of spleen were prepared by gently bursting the organ in 4 ml DMEM (Corning) in a well of a 6-well dish using the back of 3 ml syringe (Thermo Fisher). Splenocytes were passed through a 100 µm cell mesh (Corning), brought up to 25 ml with DPBS (Gibco) and spun at 300xg for 10 minutes. Erythrocytes were lysed by resuspending splenocytes in 5 ml 1x BD Pharm Lyse™ lysing solution (BD Biosciences) for 5 minutes before being brought up to 25 ml with DPBS. Cells were then pelleted at 300xg for 10 minutes, resuspended in 10 ml DPBS and counted using a Countess 3 automated cell counter (Thermo Fisher) Cells were cryopreserved in freeze media (Bambanker) for downstream cell sorting. For flow cytometry analysis, CD45+ cells were isolated from splenocytes using the EasySep™ Release Human CD45 Positive Selection Kit for Humanized Mouse Samples (Stem Cell Technologies) on an EasySep™ EasyEights Magnet (Stem Cell Technologies) according to the manufacturer's instructions. Immunophenotyping was performed using anti-human CD4-FITC (300538, RRID:AB_2562052, Biolegend), anti-human CD4-PE-Cyanine7 (300512, RRID:AB_314080, Biolegend), anti-human CD8-PE-Cyanine7 (557746, RRID:AB_396852, BD Biosciences), and anti-human CD19-FITC (302206, RRID:AB_314236, Biolegend), and cells were assessed for CAR-2A-mCherry expression using an Attune NxT flow cytometer with 96-well autosampler (Thermo Fisher). For cell sorting, cryopreserved splenocytes were thawed and stained with anti-human CD4-FITC (300538, RRID:AB_2562052, Biolegend) and anti-human CD8-FITC (557085, RRID:AB_396580, BD Biosciences) in PBS containing 1% bovine serum albumin, and an SH800S cell sorter (Sony Biotechnology) was used to sort mCherry+ and mCherry- primary human T cells that were either expressing CD4 or CD8.

Flow cytometry

Cells were stained with anti-human B2M-APC (316312, RRID:AB_10641281, Biolegend) or anti-human B2M-PE (316306, RRID:AB_492839, Biolegend) in PBS containing 1% bovine serum albumin, and an Attune NxT flow cytometer with 96-well autosampler (Thermo Fisher) was used for flow cytometry analysis. Ligand expression was confirmed for engineered HEK293T cells using anti-human CD28-PE (302907, RRID:AB_314309, Biolegend), anti-human CD20-PE (302306, RRID:AB_314254, Biolegend), anti-human CD4-PE-Cyanine7 (300512, RRID:AB_314080, Biolegend) and anti-human CD19-PE (302208, RRID:AB_314238, Biolegend). T cell immunophenotyping was performed using anti-human CD3-FITC (317306, RRID:AB_571907, Biolegend), anti-human CD4-FITC (300538, RRID:AB_2562052, Biolegend), and anti-human CD8-PE-Cyanine7 (557746, RRID:AB_396852, BD Biosciences). CD25 expression was assessed using anti-human CD25-APC (302610, RRID:AB_314280, Biolegend). Data analysis was performed using FlowJo v10 10.7.1 (FlowJo, LLC, Ashland OR).

Amplicon sequencing to assess genome editing

Next-generation sequencing was used to detect on-target genome editing in EGFP-positive, and EGFP-negative sorted HEK293T cells. Genomic DNA was extracted using QuickExtract (Lucigen) as previously described (*20*). PrimeStar GXL DNA polymerase (Takara Bio) was used to amplify the Cas9-RNP target site using the following primers: *B2M*: 5'-GCTCTTCCGATCTaagctgacagcattcgggc and 5'-GCTCTTCCGATCTgaagtcacggagcgagagag; *PDCD-1*: 5'-GCTCTTCCGATCTccgaccccacctacctaaga and 5'-GCTCTTCCGATCTgacagtttcccttccgctca. The resulting PCR products were cleaned up using magnetic SPRI beds (UC Berkeley DNA Sequencing Facility). The Innovative Genomics Institute Next-Generation Sequencing Core performed library preparation and sequencing using a MiSeq V2 Micro 2x150bp kit (Illumina). Reads were trimmed and merged (Geneious Prime, version 2022.0.1) and analyzed with CRISPResso2 (http://crispresso.pinellolab.partners.org/login).

NextSeq P2 sequencing of sorted mouse cells

Genomic DNA was extracted from sorted mCherry+ and mCherry- primary human T cells using the QIAamp DNA Mini Kit (Qiagen) according to the manufacturer's instructions. PrimeStar GXL DNA polymerase (Takara Bio) was used to amplify the *TRAC* Cas9-RNP target site using primers 5'-GCTCTTCCGATCTggggcaaagagggaaatgaga and 5'-GCTCTTCCGATCTactttgtgacacatttgtttgag. The resulting PCR products were cleaned up using magnetic SPRI beds (UC Berkeley DNA Sequencing Facility). The Innovative Genomics Institute Next-Generation Sequencing Core performed library preparation and sequencing using a NextSeq 1000/2000 P2 V3 2x150bp kit (Illumina). Reads were trimmed and merged (Geneious Prime, version 2022.0.1) and analyzed with CRISPResso2 (http://crispresso.pinellolab.partners.org/login).

TCR sequencing

Splenocytes from three Cas9-EDV-treated mice and three lentivirus-treated mice in humanized mouse experiment 2 were sorted into mCherry+ and mCherry- primary human T cells using an SH800S cell sorter (Sony Biotechnology). RNA was extracted from sorted cells using the RNeasy Mini Kit (Qiagen) according to the manufacturer's instructions. TCR a/b libraries were prepared from each sample using the SMART-Seq Human TCR (with UMIs) kit (Takara Bio) according to the manufacturer's instructions. Samples were pooled and sequenced on an Illumina MiSeq using 2×300 bp paired-end chemistry and MiSeq Reagent Kit v3 (MS-102-3003, Illumina), yielding an average sequencing depth of 1.75 million reads. FASTQ files were analyzed with Cogent NGS Immune Profiler Software (Version 1.5), using a UMI-cutoff of 3. Clonotypes were visualized using Cogent NGS Immune Viewer (Version 1.0) and a custom python script.

Immunofluorescent staining and imaging

Humanized mice were euthanized with CO2, and livers were dissected. The livers were fixed in 4% paraformaldehyde (PFA, Electron Microscopy Sciences, 15710, Hatfield, PA, USA) for 24 hours and transferred to 30% sucrose (Fisher Chemical, S5-500, Fairlawn, NJ, USA). After 3 days, the livers were embedded into cryoblocks (Tissue Plus O.C.T. Compound, Fisher HealthCare, #4548, Houston, TX, USA). 20 µm liver sections were cut with Cryostat (Cryostat, Leica, CM 3050 S, Buffalo Grove, IL, USA) and placed on microscope slides (Superfrost Plus™, Fisher Scientific, #1255015, Pittsburgh, PA, USA), and kept at -80°C until further use. The sections were blocked with a buffer including 5% normal goat serum (Sigma-Aldrich, G9023-10mL, St. Louis, MO, USA), 2% bovine serum albumin (BSA, Sigma-Aldrich, A9418-50G, St. Louis, MO, USA), 0.03% triton X-100 (Fisher Scientific, BP151-100, Eugene, OR, USA), 0.05% sodium azide (Sigma-Aldrich, 71289-5G, St. Louis, MO, USA) for 1 hour at room temperature (RT). Sections were stained for 2 hours at RT with the combination of primary antibodies as follows 1) mouse anti-β-catenin (Thermofisher Scientific, 14-2567-82, RRID:AB_1724004, Eugene, OR, USA, 1:100) and rat anti-F4/80 (Novus, NB600-404, RRID:AB_10003219, Centennial, CO, USA, 1:100); 2) mouse anti-β-catenin (Thermofisher Scientific, 14-2567-82, RRID:AB_1724004, Eugene, OR, USA, 1:100) and rabbit anti-hCD3 (Abcam, ab5690, RRID:AB_305055, Waltham, MA, USA, 1:100). Tissue sections were washed

3 times with 1xPBS (Thermofisher Scientific, 10010023, Eugene, OR, USA) and stained with the secondary antibodies for 1 hour as follows; AlexaFluor 647 goat anti-mouse (Thermofisher Scientific, A21236, RRID:AB_2535805, Eugene, OR, USA, 1:200), AlexaFluor 488 goat anti-Rat (Thermofisher Scientific, A11006, RRID:AB_2534074, Eugene, OR, USA, 1:200) and AlexaFluor 488 goat anti-rabbit (Thermofisher Scientific, A11034, RRID:AB_2576217, Eugene, OR, USA, 1:200). Tissue sections were washed again three times with 1x PBS and treated with DAPI (Sigma-Aldrich, 10236276001, St. Louis, MO, USA, 0.5 mg/mL) for 10 minutes. Then the sections were covered with cover glass slip (Micro cover glass, VWR, 48393-106, Radnor, PA, USA) with Fluoromount-G® (Southern Biotech, 0100-01, Birmingham, AL, USA). For the negative control, the sections were treated with the secondary antibodies only and DAPI (Sigma-Aldrich, 10236276001, St. Louis, MO, USA, 0.5 mg/mL).
The images of the stained livers were taken at 20X magnification with the Echo Revolve fluorescent microscope (Echo, RVL-100-G, San Diego, CA, USA) and obtained using the associated software (ECHOPro) through DAPI, FITC, Texas Red and CY5 channels.

Statistical analysis
Statistical analysis was performed using Prism v9. Statistical details for all experiments, including value and definition of N, and error bars can be found in the Figure Legends.

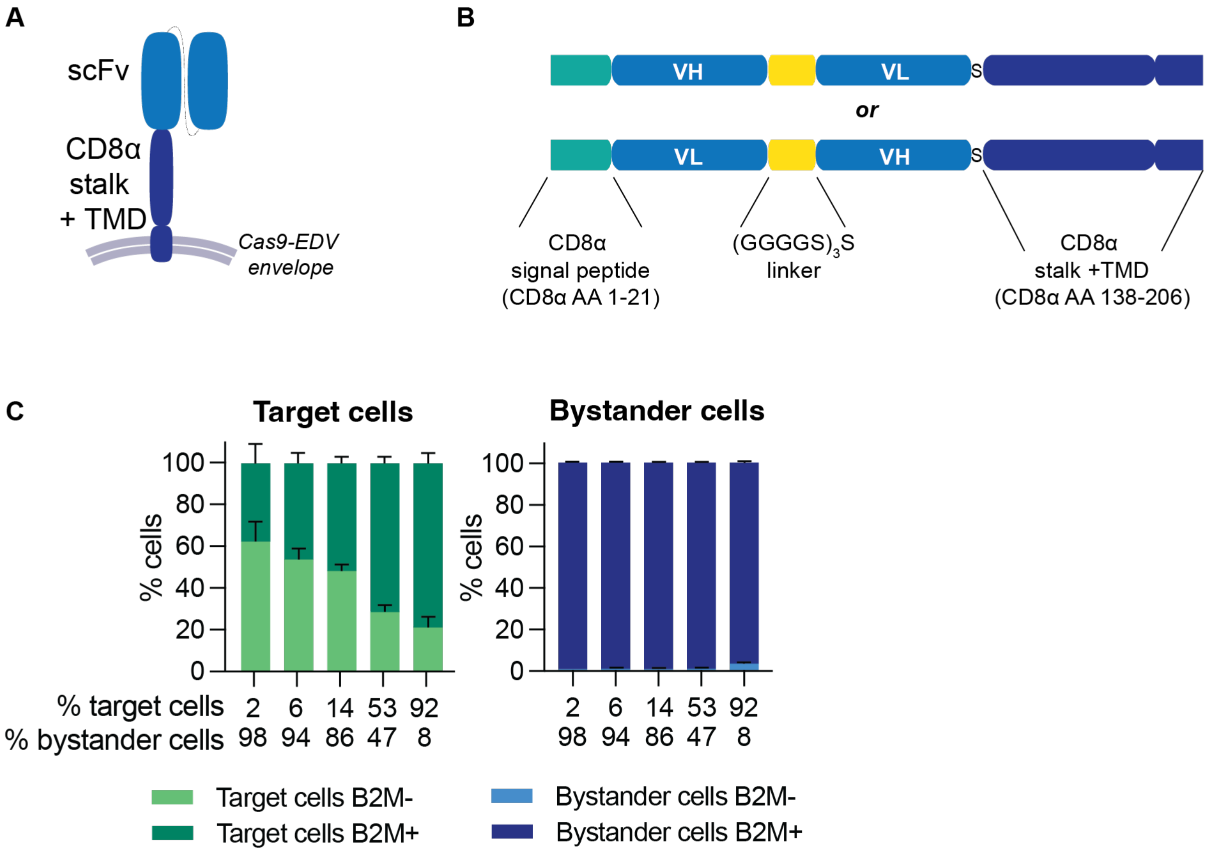


**Fig. S1. Establishing antibody-targeted Cas9-EDV delivery.** (**A**) Schematic of an antibody-derived single-chain variable fragment (scFv) targeting molecule for Cas9-EDV display. The scFv is fused to the stalk and transmembrane domain (TMD) of human CD8α. (**B**) Single-chain variable fragment (scFv) targeting molecule schematic. ScFv targeting molecules were constructed in either VH-VL or VL-VH orientations. Amino acid (AA) residues correspond to the CD8α protein sequence (UniProt P01732). TMD = transmembrane domain. S = serine. (**C**) Antibody-targeted Cas9-EDVs mediate targeted genome editing regardless of target cell frequency. CD19 EGFP HEK293T cells were mixed with HEK293T cells to achieve target cell frequencies of ~2-92%. Heterogeneous cell mixtures were challenged with antibody-targeted Cas9-EDVs (100 µl, 2.5x concentration) and *B2M* knockout was assessed in EGFP+ (on-target) and EGFP- (bystander) cells by flow cytometry 7 days post-treatment. N=3 technical replicates. Error bars represent the standard error of the mean.

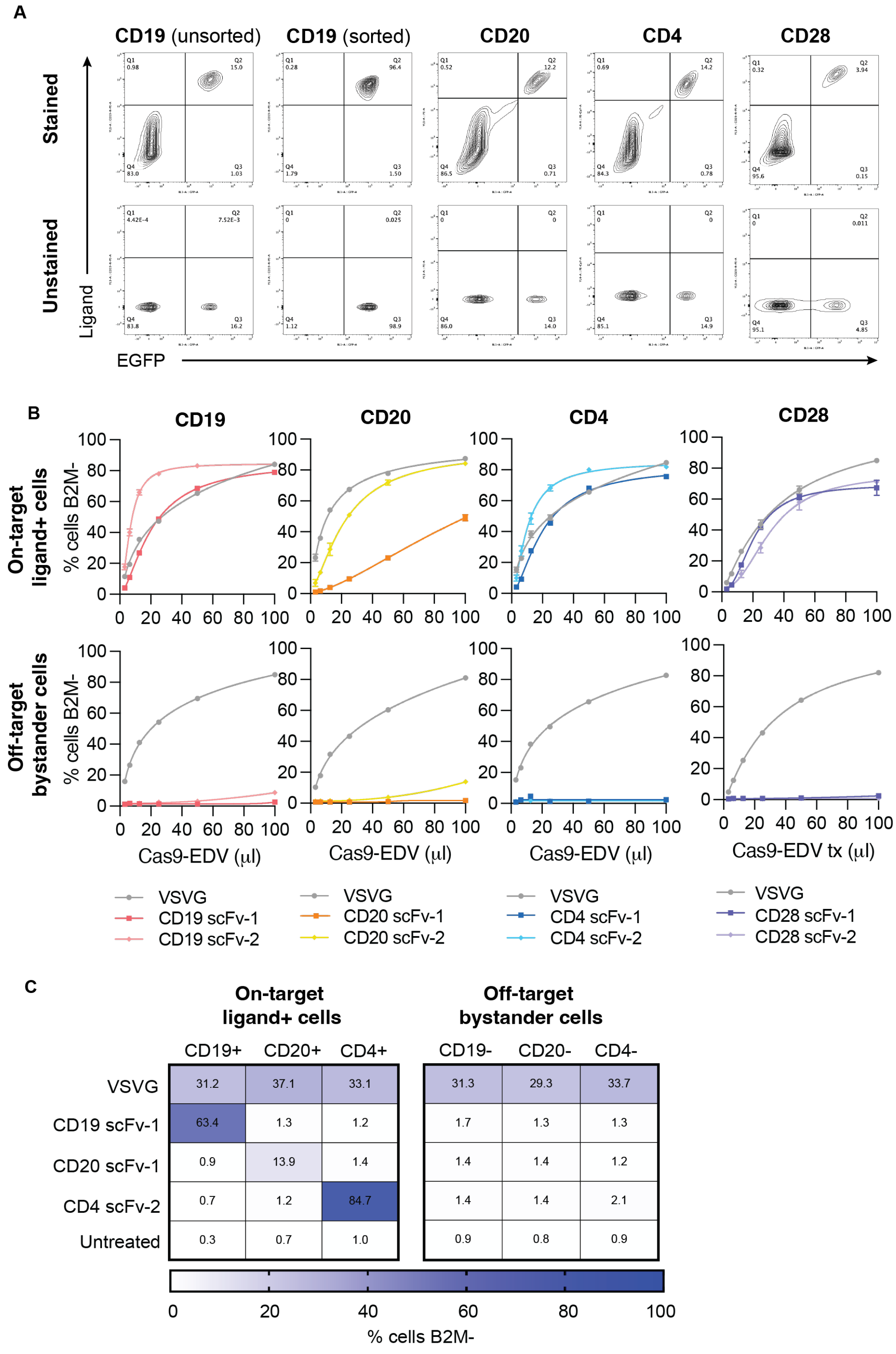


**Fig. S2. Antibody-targeted Cas9-EDVs are a programmable strategy for mediating genome editing in specific cells.** (**A**) Validation of cell-surface ligand expression in engineered cells.

Engineered 293T cells transduced to express either CD19, CD20, CD4, or CD28 and EGFP were assayed for successful ligand expression. Cells were stained with monoclonal antibodies (CD19-PE, CD20-PE CD4-PE-Cy7, CD28-PE) to verify ligand expression only in EGFP+ cells. (**B**) Various scFv targeting molecules were developed to target ligands expressed on human immune cells: CD19, CD20, CD4, and CD28. Cell-specific antibody-retargeted Cas9-EDVs activity was assessed for on-target, ligand+ cells (EGFP+) and off-target, bystander cells (EGFP-) by flow cytometry 7 days post-treatment. (**C**) Antibody-targeted Cas9-EDV-mediated genome editing is highly specific for cells expressing the scFv cognate ligand. scFv-Cas9-EDVs were tested on engineered cells expressing the cognate and noncognate ligands, and genome editing activity was measured by flow cytometry 3 days post-treatment. N = 3 technical replicates. All error bars represent the standard error of the mean.

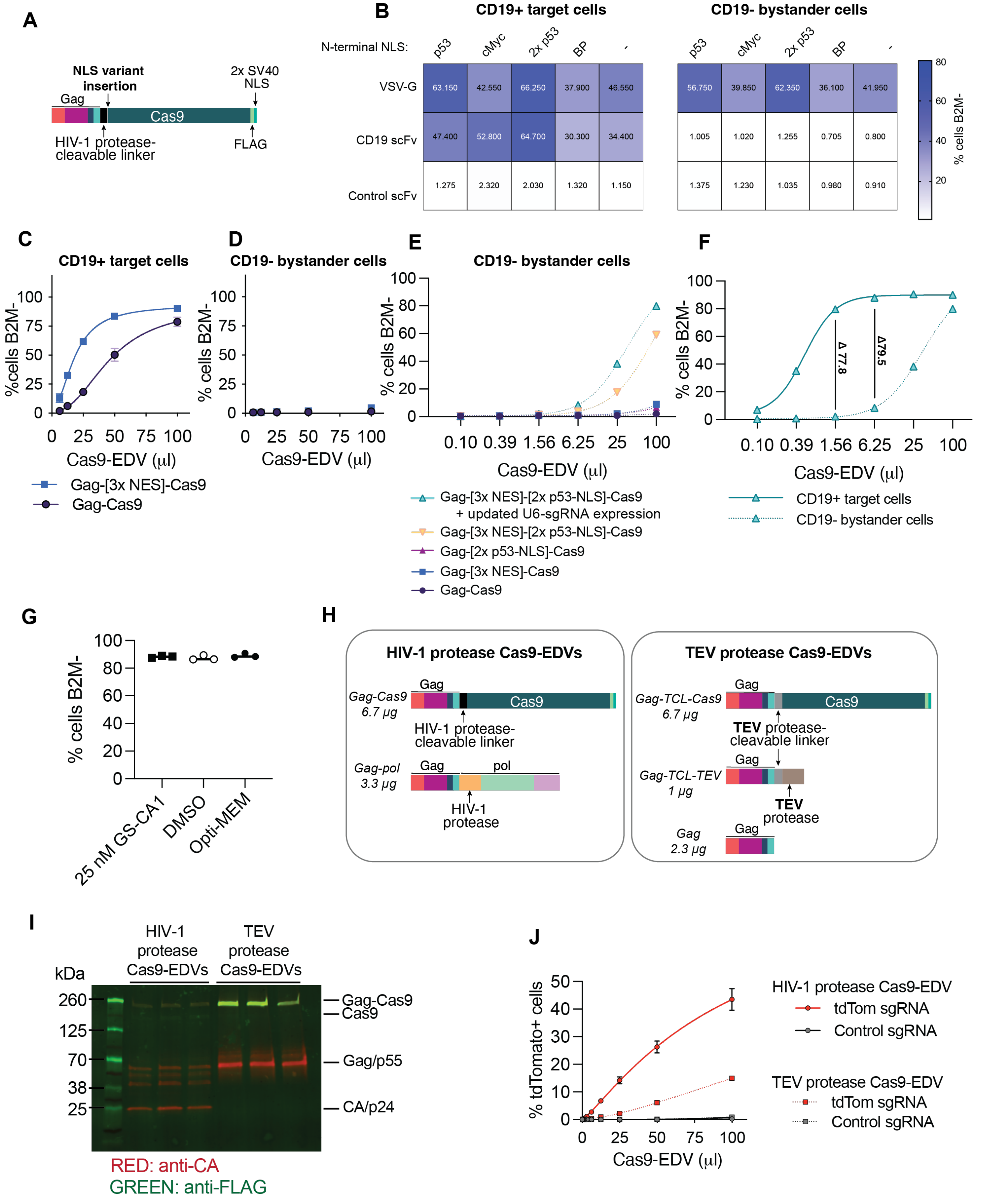


**Fig. S3. Optimization and study of Cas9-EDV activity.** (**A**) Diagram of the Gag-Cas9 plasmid indicating the site of nuclear localization signal (NLS) variant addition to the N-terminal end of Cas9. (**B**) Assessment of N-terminal Cas9 NLS additions on Cas9-EDV activity. Cas9-EDVs were used to treat a mixture of CD19+ and CD19- 293T cells. Genome editing was assessed by flow cytometry detection of B2M expression in CD19+ target cells and CD19- bystander cells 7 days post-treatment. BP = bipartite. (**C-D**) Direct comparison of genome editing in CD19+ on-target cells (**C**) and CD19- bystander cells (**D**) treated with Gag-[3x NES]-Cas9 EDVs versus Gag-Cas9 EDVs. Genome editing was assessed by flow cytometry detection of B2M expression 7 days post-treatment. (**E**) Genome editing activity comparison of CD19 antibody-targeted Cas9-EDV variants packaging *B2M*-targeted Cas9 RNPs. Expression of B2M protein was assessed by flow cytometry 7 days post-treatment in CD19-negative bystander cells (relates to Figure 3A). (**F**) Direct comparison of genome editing in on-target and bystander cells of the Gag-[3x NES]-[2x p53-NLS]-Cas9 EDV variant (relates to Figure 3A and Supplementary Figure 3E). The difference in genome-edited cells between the two cell types is indicated for 1.56 µl and 6.25 µl Cas9-EDV treatment doses. (**G**) Treatment of target cells with GS-CA1 does not alter genome editing by Cas9 RNP nucleofection. 293T cells were nucleofected with 50 pmol *B2M*-targeted Cas9 RNPs and either cultured in the presence of 25 nM GS-CA1, 0.05% DMSO, or Opti-MEM. Genome editing was assessed by flow cytometry detection of B2M expression at 3 days post-treatment. (**H**) Diagram of the Gag-TCL-Cas9 and Gag-TCL-TEV plasmids for producing TEVp-Cas9-EDVs which rely on the TEV protease to release Cas9 from Gag. TCL = TEV protease-cleavable linker. (**I**) Western blot analysis of the loss of Gag proteolytic processing in TEVp-Cas9-EDVs, as indicated by the absence of p24 and other Gag proteolytic intermediates. CA= capsid, a component of the gag polypeptide. (**J**) Genome editing activity of TEVp-Cas9-EDVs packaging Cas9 RNPs that activate tdTomato expression upon genome editing of a murine neural progenitor cell line harboring a loxP-STOP-loxP-tdTomato reporter (*54*). Genome editing was assessed by flow cytometry detection of tdTomato reporter activation 7 days post-treatment. N=3 technical replicates were used in all experiments.

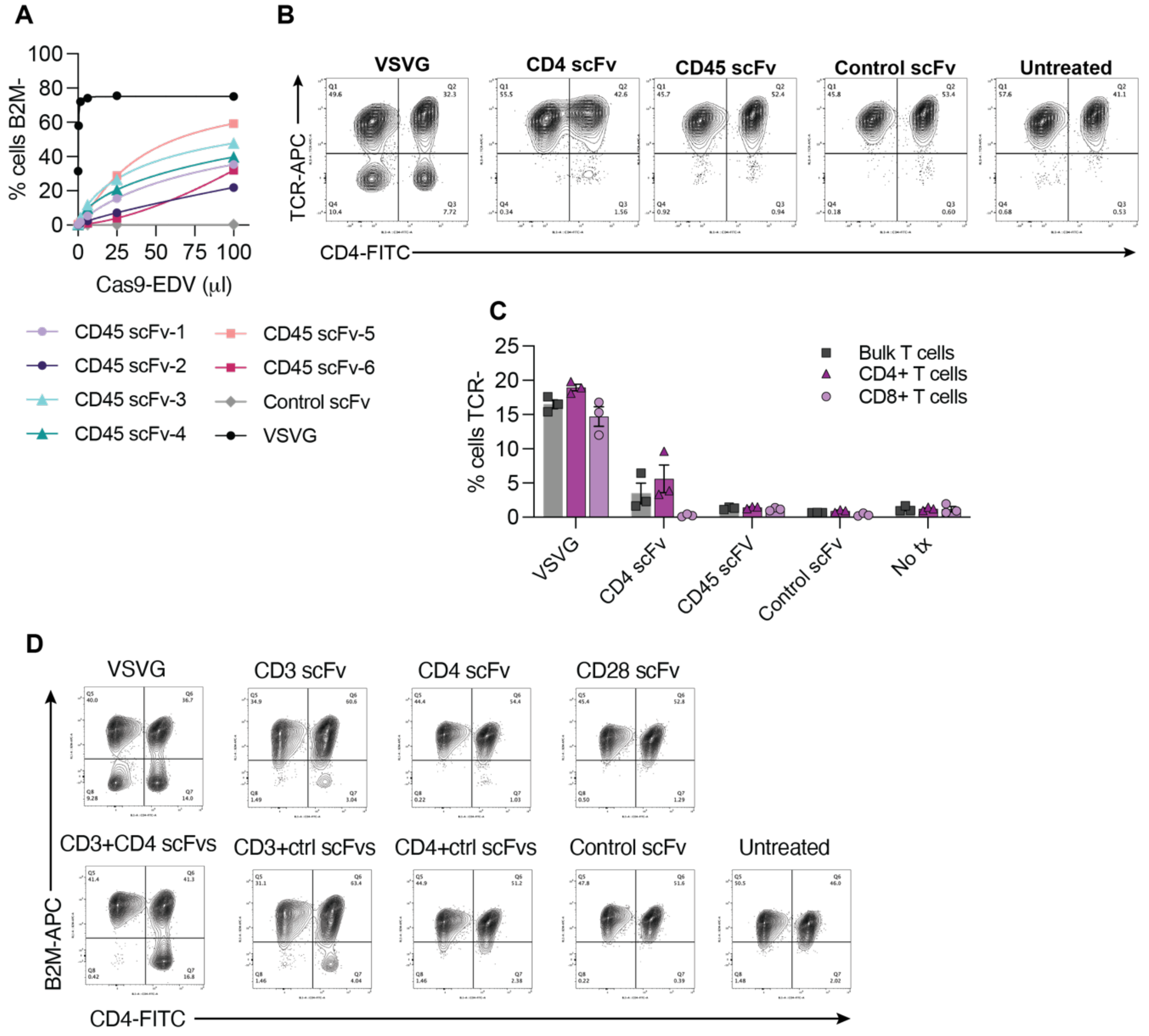


**Fig. S4. Assessment of T cell-targeting molecules for Cas9-EDV delivery.** (**A**) Assessment of human CD45 targeting molecules. A panel of CD45-targeted Cas9-EDVs packaging *B2M*-targeted Cas9 RNPs were produced and concentrated 13.8-fold. 100 µl was used to treat 30k Jurkat cells, and B2M expression was assessed by flow cytometry on day 3. (**B**) Assessment of CD4 and CD45-targeted Cas9-EDVs in activated primary human T cells. Cas9-EDVs packaging *TRAC*-targeted Cas9 RNPs were produced and 1.15x10$^{7}$ Cas9-EDVs were used to treat 15k activated T cells. T cell receptor (TCR) loss, indicative of *TRAC* genome editing, was assessed by flow cytometry 5 days post-treatment. Representative plots are shown, and results are quantified in (**C**). CD4 scFv-1 and CD45 scFv-5 targeting molecules were used. (**D**) Assessment of multiplexing targeting molecules for delivery to activated primary human T cells. 15k activated T cells were treated with 1.38x10$^{8}$ Cas9-EDVs packaging *B2M*-targeted Cas9 RNPs, and flow cytometry analysis of B2M expression was assessed on day 6. Representative flow cytometry plots are shown.CD3 scFv-3, CD4 scFv-2, and CD28 scFv-2 targeting molecules were used. Error bars represent the standard error of the mean. N=3 technical replicates were assessed (**A & C**).

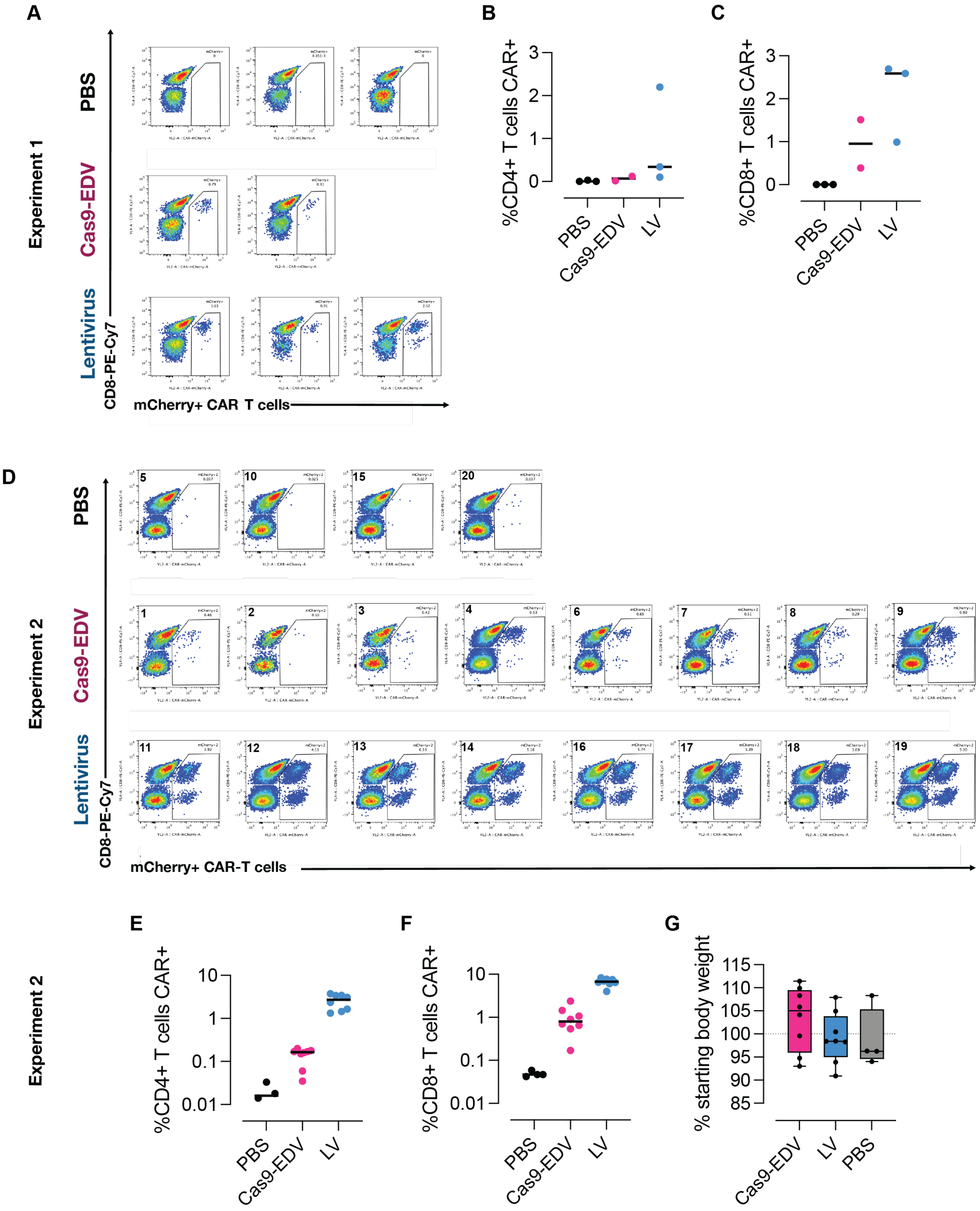


**Fig. S5. Assessment of CAR T generation *in vivo* using T cell-targeted vectors.** (**A**) Analysis of mCherry+ CAR T cells in spleens from humanized mouse experiment 1. Mice were treated with Cas9-EDV, lentivirus (LV), or PBS and flow cytometry analysis was performed 10 days post-treatment. Flow cytometry quantification of CD4+ (**B**) and CD8+ CAR T cells (**C**). (**D**) Analysis of mCherry+ CAR T cells in spleens from humanized mouse experiment 2. Mice were treated with Cas9-EDV, lentivirus (LV), or PBS and flow cytometry analysis was performed 10 days post-treatment. Upper left quadrant numbers are mouse identifiers. Flow cytometry quantification of CD4+ (**E**) and CD8+ CAR T cells (**F**). (**G**) Quantification of body weight change in humanized mice from experiment 2. The dotted line indicates 100%. Black lines indicate the median (**B-C, E-G**).

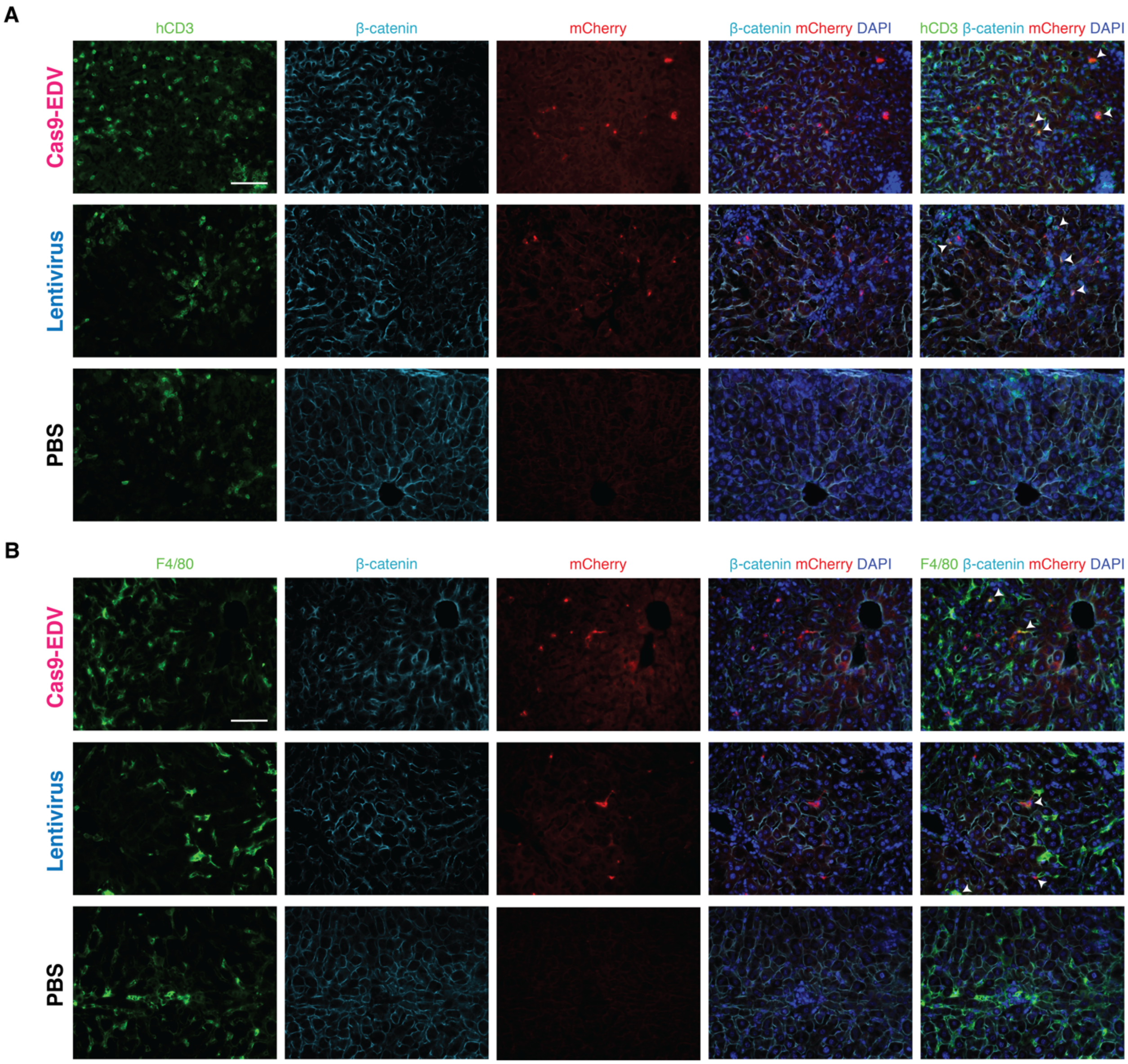


**Fig. S6. Analysis of delivery to off-target cells *in vivo*.** (**A-B**) Representative confocal microscopy images of the livers from PBMC-humanized mice 10 days post systemic administration of T-cell targeting Cas9-EDV and lentivirus demonstrating that mCherry+ (red) cells co-localize with human CD3+ (green) T cells (**A**) and F4/80+ (green) phagocytic cells (**B**) but not β-catenin+ (teal) hepatocytes. Scale bar, 65 μm.

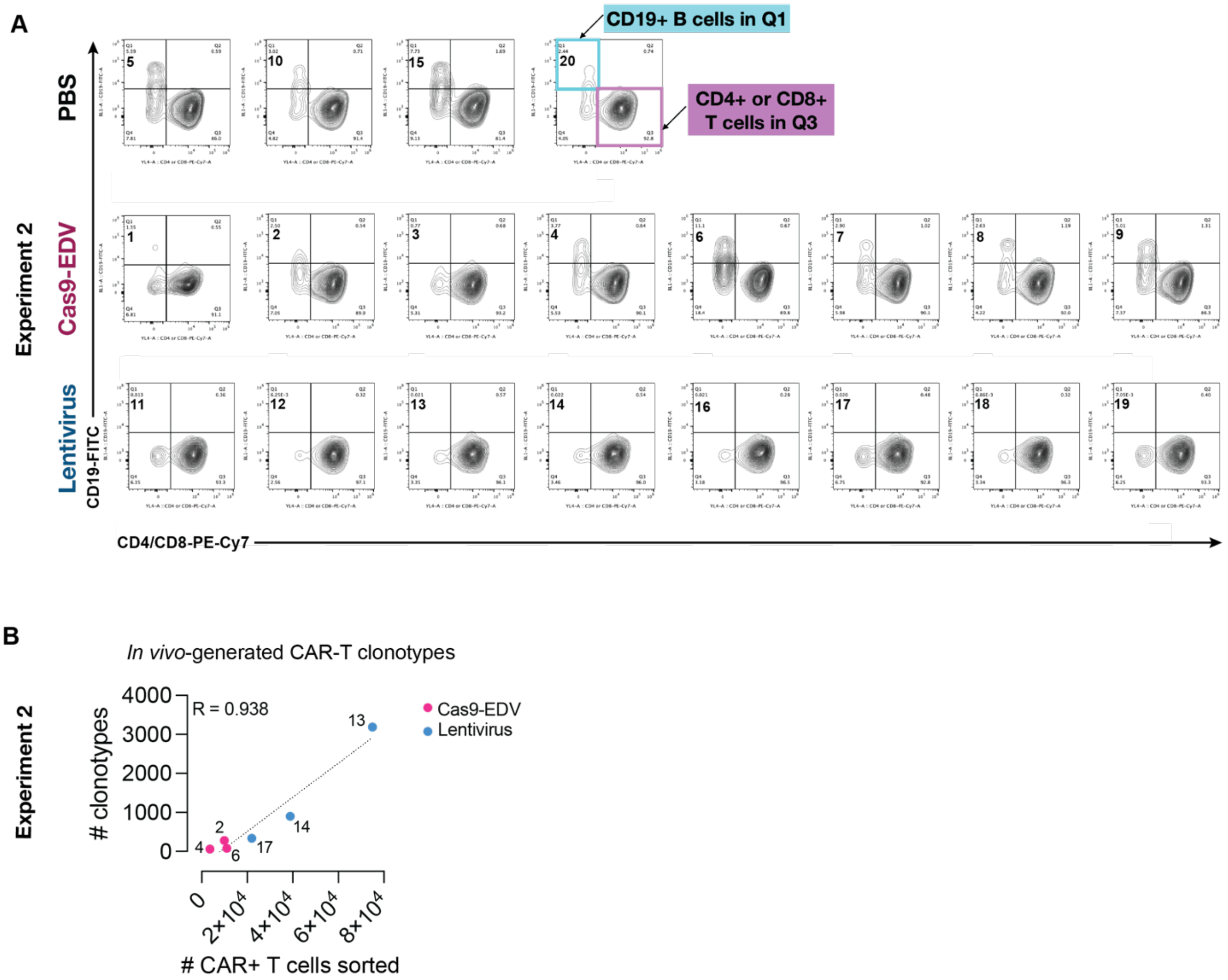


**Fig. S7. Assessment of CD19+ B cells and CAR T cell receptor clonality following *in vivo* cellular engineering.** (**A**) Flow cytometry analysis of CD19+ B cells in Cas9-EDV- or lentivirus-treated mice 10 days post-administration. Human CD45+ cells were isolated from humanized mouse spleens prior to flow cytometry analysis. Upper left quadrant numbers are mouse identifiers. (**B**) Correlation between the number of *in vivo*-generated CAR T clonotypes versus the number of CAR T cells sorted (Pearson's correlation coefficient, R = 0.938). Point labels indicate the unique mouse analyzed.
